# Allelic variation at tRNA genes in three nematode species indicates mutation load despite strong purifying selection

**DOI:** 10.1101/2025.10.14.682033

**Authors:** Avery Davis Bell, Corinne Simonti, Hector Baños, Ling Wang, Christine Heitsch, Joseph Lachance, Annalise B. Paaby

## Abstract

Cytosolic transfer RNAs, which are encoded as hundreds of genes in nuclear genomes, experience exceptionally high mutation rates and have been hypothesized to confer substantial mutational load in natural populations. Although this phenomenon appears universal across multicellular eukaryotes, a comprehensive characterization of standing variation in tRNA repertoires is still lacking in any system. Here, we resolve within-species allelic variation in nuclear-encoded tRNAs in three nematode species: *Caenorhabditis elegans*, *C. briggsae*, and *C. tropicalis*. We show that these genes carry signatures of high rates of historical transcription-associated mutagenesis and of purifying selection, resulting in allelic variation that includes pervasive instances of within-gene mismatches between the amino acid recognized by the tRNA backbone and that indicated by the anticodon. Furthermore, patterns of tRNA genomic organization and variation differ markedly from those of protein-coding regions. Individual genomes harbor distinct complements of tRNA genes with predicted functional differences, an observation that coincides with recent evidence that variation in tRNA expression and regulation contributes to human disease. Our findings offer an entry point for identifying the micro-evolutionary processes that act on tRNA repertoires, and in turn connecting those processes to the macro-evolutionary patterns that have more frequently been the focus of study.

**Significance statement:** Transfer RNAs (tRNAs) are ancient and essential molecules that deliver amino acids to the growing polypeptide chain during protein synthesis. Despite their importance, little is known about how the genes encoding tRNAs vary among individuals, even though they accumulate large numbers of mutations due to their high rates of expression. Here, we report that three nematode species, including the model organism *C. elegans*, show within-species variation in their tRNA gene repertoires, which may cause differences in protein synthesis across individuals. Our findings may represent patterns common across multicellular species and they set the stage for future investigation into how this diversity influences cellular function and organismal fitness.

## Introduction

Transfer RNAs are essential molecules that deliver individual amino acids to the growing polypeptide chain during protein synthesis. Universal across the tree of life, tRNAs are encoded as genes in nuclear and organelle genomes and exhibit deep conservation of the stereotypical ‘cloverleaf’ secondary structure (**Figure 1A**) and L-shaped tertiary structure integral to their molecular function. tRNAs are specific to their cognate amino acid, typically occurring as one of 22 isotypes (corresponding to the 20 canonical amino acids plus two modified amino acids). The isotype of a tRNA is determined by its anticodon, which recognizes codons in the mRNA, as well as other sequence identity elements necessary for binding specificity of the appropriate aminoacyl-tRNA synthetase; these may reside in the acceptor stem, for example (**Figure 1A**) (Giege and Eriani 2023).

**Figure 1.**
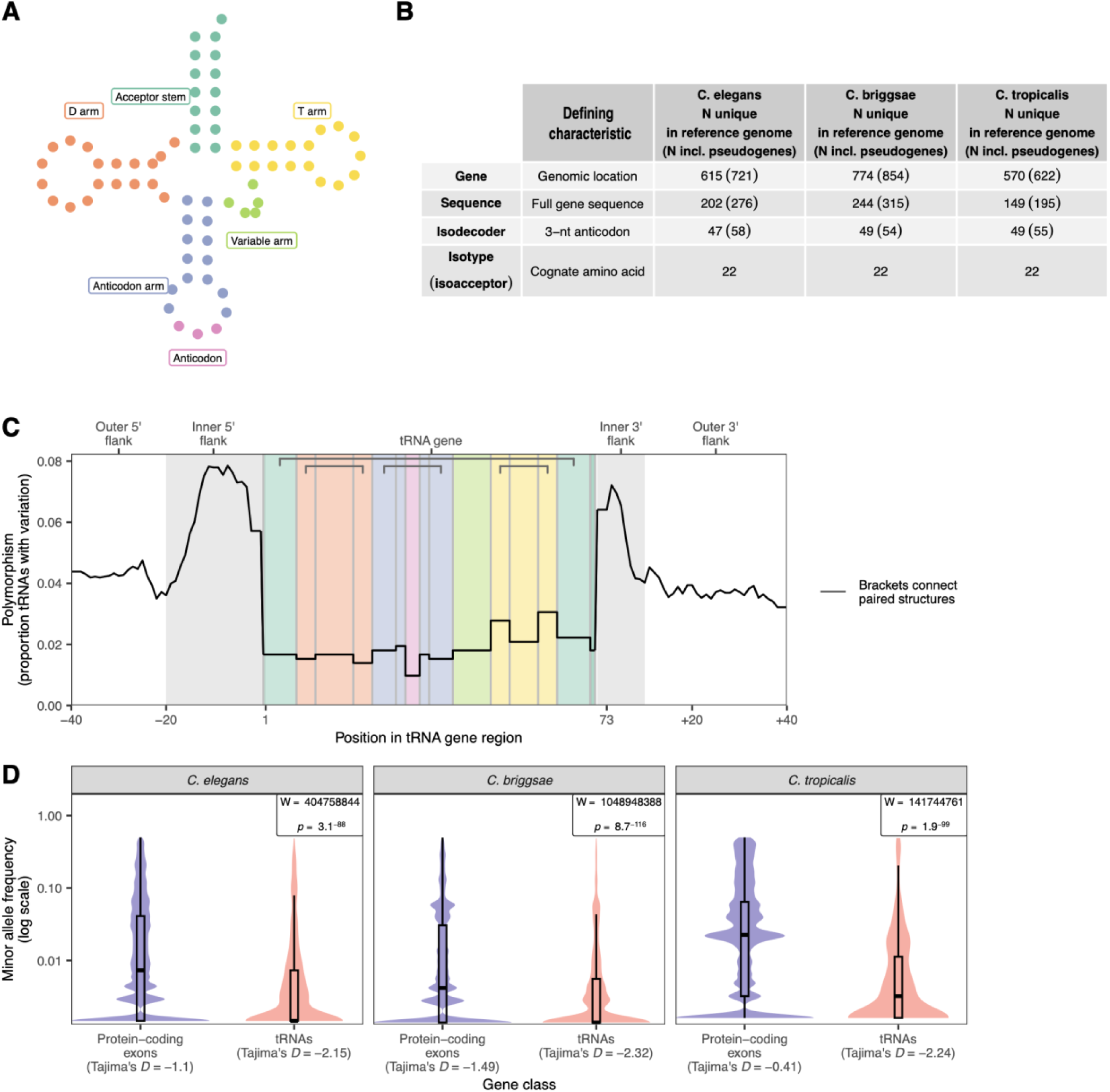
Structural, functional, sequence similarity, and transcription-associated mutagenesis (TAM) of tRNA genes in *Caenorhabditis* nematodes. **A,** “Cloverleaf” secondary structure of a canonical 72 bp tRNA, with each base plotted as a circle and colored and labeled by the component of the tRNA structure it is part of (linker bases included with the neighboring arm). **B,** The number of tRNA genes in the reference genomes of three *Caenorhabditis* species: individual genes (which have distinct genomic locations), distinct full-tRNA sequences (which may occur at multiple genomic locations), distinct anticodons (isodecoders), and distinct amino acids (isotypes/isoacceptors). Numbers outside of parentheses represent putatively functional tRNA genes, while numbers in parenthesis include these as well as predicted non-functional tRNA genes. **C**, Variation across tRNA gene regions in *C. elegans* for all tRNA genes in the reference genome combined (n = 720, includes all tRNAscan-SE predicted tRNAs including pseudogenes): the y axis is the proportion of tRNAs with mutational variation (variants with minor allele frequency < 0.05) observed in the 687-strain population at that base (5 bp moving window average). Regions in the tRNA gene itself are color coded according to secondary structure (as in **A)** while flanking regions showing the influence of TAM without selection are in white and gray. (Linker bases excluded.) **D,** Minor allele frequency (MAF) spectrum of variants (SNVs and INDELs) in protein-coding exons and putatively functional tRNA genes; between-gene set Mann-Whitney test results (top right) and Tajima’s *D* values are annotated. Boxplots overlaid to show median (bold center bar), 25^th^ and 75^th^ data percentiles (box), and 1.5x interquartile range (whiskers). *N* variants per class per species in Table S3.

However, despite ancient conservation of this molecular structure-function relationship, tRNA sequences in nuclear genomes also undergo extensive evolutionary turnover. These cytosolic tRNAs typically occur as hundreds of genes in eukaryotic genomes, often including dozens per isotype (Bermudez-Santana et al. 2010; Goodenbour and Pan 2006). Across eukaryotic lineages, cytosolic tRNA repertoires exhibit copy number variation, reassignment of tRNA identity from one cognate amino acid to another (isotype switching), and frequent gains and losses, including by pseudogenization following accumulation of mutations (Bermudez-Santana et al. 2010; Rogers et al. 2010; Iben and Maraia 2012; Wang and Ruvinsky 2012; Rogers and Griffiths-Jones 2014; Velandia-Huerto et al. 2016). The fraction of tRNA loci that share syntenic orthology among extant primate species is estimated to be between one third and one half (Velandia-Huerto et al. 2016); between the species *Caenorhabditis elegans* and *C. briggsae*, also about one half (Wang and Ruvinsky 2012).

The evolutionary dynamism of tRNAs is likely driven by the strong, opposing evolutionary forces they experience. tRNAs are among the most highly transcribed genes in the genome and consequently are subject to extremely high rates of transcription-associated mutagenesis (TAM) (Thornlow et al. 2018). Like other aspects of tRNAs, this phenomenon appears universal: polymorphism at tRNA loci in humans, *Mus musculus*, *Drosophila melanogaster*, and *Arabidopsis thaliana* show strong signatures of TAM (Thornlow et al. 2018), which induces an excess of C to T and G to A mutations on the coding strand (Green et al. 2003; Jinks-Robertson and Bhagwat 2014). During tRNA transcription, flanking regions just upstream and downstream of the gene are likewise subject to TAM. These flanking regions exhibit dramatically elevated rates of substitution relative to the gene body, providing a record of historical exposure to TAM and indicating that functional tRNA sequences are subject to strong purifying selection (Thornlow et al. 2018). Modeling the TAM signature via a sequence substitution model, we find that standing variation in *C. elegans* tRNAs is well explained by historical exposure to TAM (Banos et al. 2025).

Thornlow and colleagues (2018) have proposed that the elevated rate of TAM on nuclear-encoded tRNAs may confer a significant mutation load on eukaryotic genomes. Exploration of this hypothesis ultimately requires evaluating the fitness consequences of standing genetic variation in tRNAs within extant populations. Mutations to mitochondrial tRNAs have long been known to cause an array of diseases (Florentz et al. 2003; Goto et al. 1990; Kobayashi et al. 1990; Suzuki et al. 2011; Yarham et al. 2010), but the occurrence of mt-tRNAs as typically single-copy genes makes them both more vulnerable to impacts of mutation and more tractable to study. At nuclear-encoded tRNAs, tRNA gene redundancies make it difficult to identify, and empirically test, variants with functional consequences, though mutations in human cytosolic tRNAs predicted to cause amino acid mis-loading can induce biochemical consequences (Davey-Young et al. 2024; Tennakoon et al. 2025). However, recent studies indicate growing interest in allelic variation of cytosolic tRNAs, especially in the context of disease (Schoenmakers et al. 2016; Thornlow et al. 2018; Lant et al. 2019; Pinzaru and Tavazoie 2023; Murillo-Recio et al. 2025). In humans, studies with small sample sizes reveal substantial variation at tRNA genes, even as they likely under-sample the breadth of variation (Iben and Maraia 2014; Darrow and Chadwick 2014; Berg et al. 2019). Larger-scale, lower-coverage sequencing also indicates extensive variation (Parisien et al. 2013; Lant et al. 2019), though these approaches do not capture the rare variants that likely contribute disproportionately to functional differences. The consensus view holds that the full extent of cytosolic tRNA variation remains underestimated (Lant et al. 2019).

In parallel, the rapidly emerging field of tRNA regulation has demonstrated a link between the dysregulation of cytosolic tRNAs and disease. These findings, facilitated by new approaches to analyzing tRNA expression, show that tRNA regulatory control is complex and dynamic across age, cell type, and environmental cues (Pinzaru and Tavazoie 2023; Suzuki 2021). These studies directly connect cytosolic tRNA activity to organismal fitness, though so far this research area has largely overlooked the consequence of mutational variation on post-transcriptional tRNA function.

To begin to evaluate the fitness consequences of mutational variation in cytosolic tRNAs, we first require comprehensive characterization of tRNA allelic variation at the population level. The universality of the genetic code, deep conservation of translational machinery, and ubiquity of TAM-induced substitutions at tRNA loci (Thornlow et al. 2018) indicate that insights from any system should be informative across eukaryotes. For example, despite their differences in organismal complexity, the genomes of flies and worms to humans and chimps encode comparable number of tRNAs (Goodenbour and Pan 2006), which vary by isotype copy number and reflect adaptation to codon usage (Duret 2000; Qian et al. 2012). Here, we evaluate genotype data for hundreds of wild strains in three species of *Caenorhabditis* nematode worms: the model organism *C. elegans* and related species *C. briggsae* and *C. tropicalis* (Crombie et al. 2024). These species diverged from each other ∼10-25 million years ago, exhibit ∼20-30% synonymous site differences (Stevens et al. 2022; Fusca et al. 2025), and are androdioecious, *i.e.*, hermaphroditic and largely self-fertilizing in nature. This mating system may permit the persistence of deleterious variants due to reduced recombination and less efficient selection (Barriere and Felix 2005a; Dolgin et al. 2007; Loewe and Cutter 2008; Frézal and Félix 2015; Fusca et al. 2025), enabling the detection of rare variants and alleles.

In this study, we resolved within-species allelic variation in nuclear-encoded tRNAs from >620 wild strains in each of the three *Caenorhabditis* species. Then, we asked (i) what evolutionary forces shape these tRNA repertoires, (ii) to what extent the observed allelic variation may affect tRNA function, including via within-gene isotype mismatches between the tRNA backbone (scaffold) and the anticodon, and (iii) whether the genomic organization of tRNA genes departs from that of protein-coding regions and that predicted by genome architecture.

## Results

### tRNA repertoires in *Caenorhabditis*

To characterize the pool of tRNA genes encoded in the nuclear genomes of *C. elegans*, *C. briggsae*, and *C. tropicalis*, we first computationally predicted tRNA-like sequences in the three reference genomes. As tRNAs exhibit highly conserved ‘cloverleaf’ secondary structure and corresponding genomic sequence motifs (**Figure 1A**), putatively functional tRNAs and non-functional tRNA-like sequences (pseudogenes) can be identified with high confidence from sequence data (Chan et al. 2021). Each tRNA gene belongs to multiple nested classes (**Figure 1B**): the gene itself defined by its unique location, a set of one or more genes with identical sequence, an isodecoder class of genes that share the same anticodon and therefore cognate amino acid, and an isotype (also called isoacceptor) class that shares the same cognate amino acid but not necessarily the same anticodon. Each species encoded tRNAs for all 20 canonical amino acids as well as initiation-specific methionine (iMet) and selenocysteine (SeC); the total number of putatively functional tRNA genes within each reference genome was 615 (*C. elegans*), 774 (*C. briggsae*), and 570 (*C. tropicalis*) (**Figure 1B**). Within species, these genes often shared sequence identity, such that the number of unique tRNA sequences ranged from 149-244 per reference genome, in turn comprised of 48-49 unique anticodon sequences (**Figure 1B, Table S1-2**). These patterns were consistent with known tRNA gene complements (Chan and Lowe 2009, 2016).

We first investigated isotype and isodecoder frequency at tRNA genes. While copy number varied substantially, with some isotypes differing by a third or more across species (**Table S1**), overall isotype frequencies were broadly similar (Pearson’s *R^2^* ≥ 0.92, *p* < 4.3 x 10^-13^ for gene number across isotypes). For example, glycine and leucine rank first and second in isotype prevalence in *C. elegans* and *C. briggsae*; in *C. tropicalis*, they rank third and first, with proline the second most prevalent isotype. These differences represent relative isotype representation shifts of up to 30% across species, though the individual isotype frequencies are modest within the full repertoire (*e.g.*, a shift in proline frequency from 0.068 in *C. elegans* to 0.089 in *C. tropicalis*). In all three species, selenocysteine (SeC) was the rarest tRNA encoded amino acid (**Table S1**), consistent with the rarity and unique role of SeC across the tree of life (Chan and Lowe 2016; Commans and Bock 1999).

At the codon level, most codon sequences (corresponding to the anticodon of the transcribed tRNA molecule) appeared in multiple tRNA genes within each reference genome. The most frequently observed codons were TCC/glycine in *C. elegans* and *C. briggsae* (36 and 60 genes, respectively) and TGG/proline in *C. tropicalis* (38 genes) (**Table S2**). Each reference genome contained tRNA genes with a single instance of a given codon (three in *C. elegans*, five in *C. briggsae*, two in *C. tropicalis*), though these were not necessarily the same codons, and several (out of 61 possible) were entirely absent. Missing isodecoders are a persistent feature of tRNA repertoires in all living systems (Chan and Lowe 2016), even as the codons themselves appear in protein-coding genes throughout the genome; wobble pairing at the third position permits codon reading by non-cognate tRNAs (Crick 1966; Roth 2012). Missing isodecoders tended to be consistent across the three species: of the 18 codon sequences missing in any species, 13 were missing in all three (**Table S2**).

### *Caenorhabditis* tRNA genes exhibit signatures of transcription-associated mutagenesis

tRNAs in humans, mice, *Arabidopsis*, and *Drosophila* show evidence of exceptionally high rates of mutation, incurred because of their high rate of transcription and evidenced by patterns of intraspecific polymorphism (Thornlow et al. 2018). To assess the role of transcription-associated mutagenesis (TAM) in our system, we examined population variation at tRNA loci and their flanking genomic regions in our three species. During transcription, a gene and its flanking regions are unwound and the unbound, non-template strand is vulnerable to mutation. For tRNAs, the elevated rate of transcription induces an exceptionally high mutation rate; a substantial fraction of the mutations are purged from the gene body by purifying selection, but not from the flanking regions, which provide a historical record of mutagenic activity (Jinks-Robertson and Bhagwat 2014; Thornlow et al. 2018). Consistent with the reports in other systems (Thornlow et al. 2018) and our earlier findings in *C. elegans* (Banos et al. 2025), tRNA genes and their flanking regions exhibited signatures of TAM in the genetic variation of wild strains in all three species (**Figure 1C, S1, S2**).

Specifically, tRNA gene flanking regions had exceptionally high mutational variation, with up to 7.6-12.9% of tRNA genes exhibiting mutations at a given base across species (**Figure 1C, Figure S1**). The substitutions most often induced by TAM, G>A and C>T (Gaillard and Aguilera 2016; Jinks-Robertson and Bhagwat 2014), tended to be the most frequent mutations in both upstream tRNA gene flanking regions and within tRNA gene bodies (**Figure S2**). Consistent with strong purifying selection, tRNA gene bodies exhibited many fewer mutations than flanking regions (**Figure S1**). Nevertheless, we observed mutations across the whole length of the tRNA in each of the three species (**Figure S1**). As we observed in a *C. elegans*-specific analysis (Banos et al. 2025), polymorphism was elevated at paired stem sites relative to loops. This is likely driven by increased susceptibility to TAM arising from the higher GC content at paired sites (Banos et al. 2025), though it is also consistent with stronger purifying selection on loops, which indicate greater fitness costs when mutated (Li et al. 2016).

Under the assumption that the rate of deleterious mutation and subsequent purifying selection are exceptionally high for tRNAs, we hypothesized that tRNA genes might be enriched for low frequency variants relative to other functionally important elements. Indeed, compared to protein-coding genes, variants in putatively functional tRNA genes tended to have lower minor allele frequencies (**Figure 1D**), and these variants were much more likely to be present in the population as singletons (49-53% singleton variants in tRNAs vs. 24-37% in protein-coding regions across species, **Table S3**). This excess of rare alleles was likewise reflected in lower estimates of Tajima’s *D* (**Figure 1D, Table S4**), which assesses site frequency spectra by comparing pairwise differences to the number of segregating sites (Tajima 1989). The difference between protein-coding genes and tRNAs remained strongly significant for each species comparison when all tRNAs, including those predicted to be non-functional, were included, though non-functional sequences had higher minor allele frequencies and smaller singleton proportions (**Table S3**). This shift likely reflects stronger purifying selection on active tRNA loci.

### Allelic variation at *Caenorhabditis* tRNA genes with putatively functional consequences

Next, we sought to characterize intraspecific allelic variation at tRNA genes within the three *Caenorhabditis* species. To do this, we reconstructed full-length alleles for each gene in each wild strain (>620 per species), using variant genotype data from CaeNDR (Crombie et al. 2024) and the gene sets from the reference genomes. Some tRNA genes were invariant, exhibiting only one allele; variable genes harbored up to 17 alleles across the populations (**Figure 2, Figure S3**). Among putatively functional tRNA genes, 38.8-83.3% showed allelic variation across the three species (**Figure S3**). Across all nuclear-encoded tRNAs, including putative pseudogenes predicted to be non-functional, individual tRNA isotypes comprised 1-139 alleles in *C. elegans* (of 3574 total unique alleles at all tRNA genes in the entire population), 4-328 in *C. briggsae* (of 3574 total), and 1-99 in *C. tropicalis* (of 1048 total).

**Figure 2.**
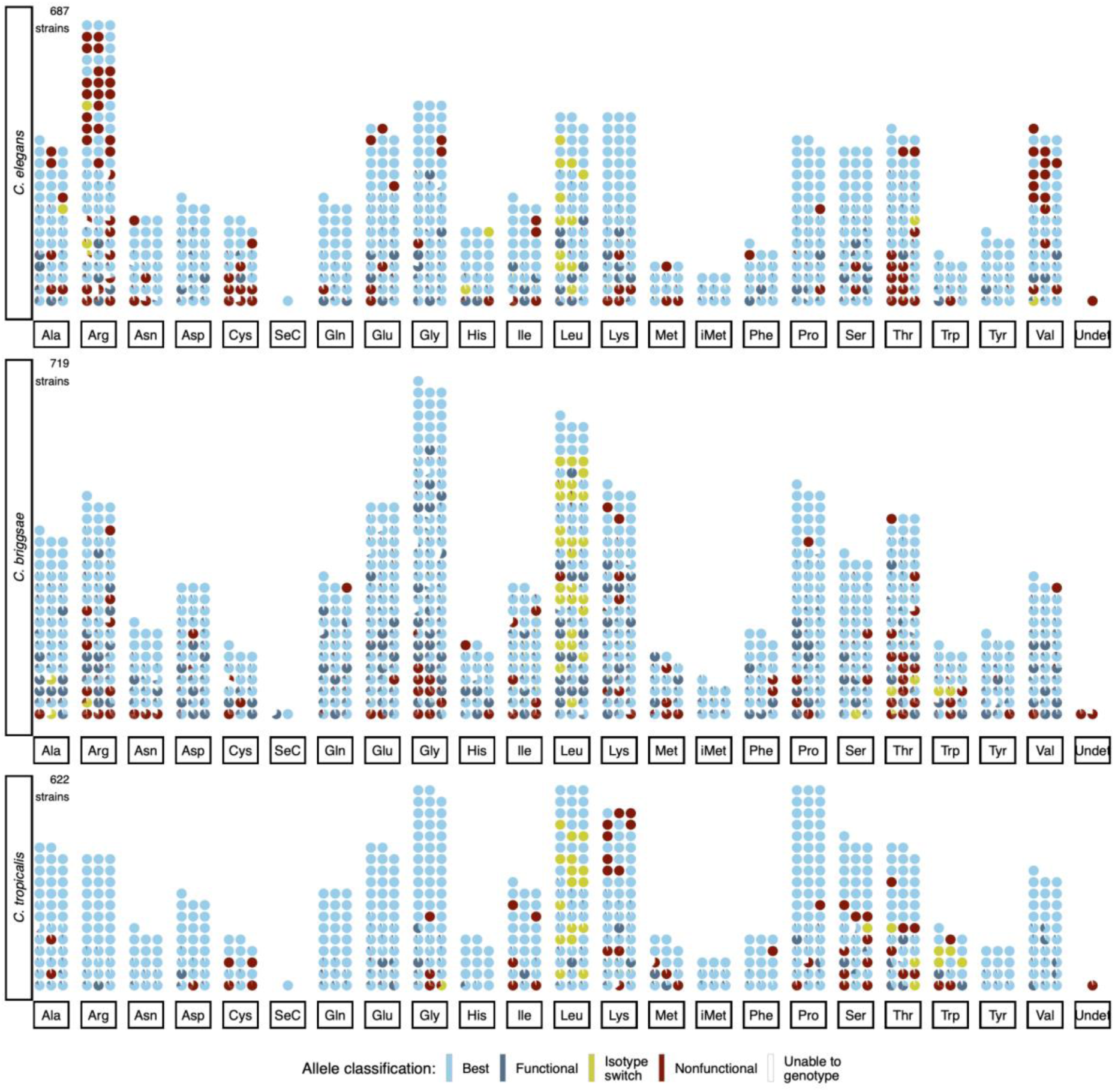
Allelic diversity at all tRNA genes in *Caenorhabditis* reference genomes. Each tRNA gene in the reference genome of each species is represented by a circle, grouped by the amino acid (isotype) predicted to match its backbone; slices of the circle correspond to different alleles of that gene (alleles with frequency < 0.05 plotted as having frequency = 0.05). Allele slices are colored by inferred functional classification (main text); white slices correspond to strains where one or more variants in that gene had missing genotypes. Genes may have multiple alleles of the same classification (color), but may have one and only one ‘best’ allele. The number of strains for which allelic data was generated is in the top left corner. Details on each gene, including zooms of each pie chart, are available via a searchable web app at https://wildworm.biosci.gatech.edu/trnarefalleles/.

The extensive allelic diversity over a large number of tRNA genes resulted in a unique complement of putatively functional tRNA alleles for each strain of *C. elegans* and *C. briggsae*. In *C. tropicalis*, which has lower average genetic diversity (Noble et al. 2021), 87% of strains (545 of 622) exhibited unique tRNA allelic complements; the remaining strains shared tRNA allele identity in sets of two or three strains, with the exception of one allelic combination, which was shared across nine strains.

To investigate potential functional relevance of this allelic variation, we developed allele classifications by combining information about: the predicted amino acid for that tRNA, inferred from the tRNA backbone sequence excluding the anticodon; the amino acid matching the anticodon; and the estimated tRNA folding score (all from tRNAscan-SE). Alleles for which the anticodon and backbone predicted the same amino acid were classified as ‘Best’ if they had the highest RNA folding score or ‘Functional’ if they had a putatively functional score lower than that of the ‘Best’ allele; in practice, any of the ‘Functional’ alleles might truly be most functional. Alleles were classified as ‘Nonfunctional’ by low folding scores or non-identifiable anticodons, and ‘Isotype switch’ by a mismatch between the amino acid predicted from the backbone sequence and the amino acid matching the anticodon; isotype switching has long been observed across tRNA evolution at interspecies timescales (Rogers and Griffiths-Jones 2014).

Within each species, a substantial proportion of the nuclear tRNA genes exhibited putatively functional allelic diversity, with alleles belonging to all categories (**Figure 2**). Of the genes classified as ‘Nonfunctional’ in the reference genomes (**Figure 1B**), >91% in each species exhibited only ‘Nonfunctional’ alleles (or missing data) across the strains. However, genes in all three species harbor both functional and nonfunctional alleles (46 in *C. elegans*, 99 in *C. briggsae*, and 10 in *C. tropicalis*), suggesting that some strains in the population express viable tRNAs for that specific gene while others do not (**Figure 2**). tRNA isotypes varied in the degree of predicted functional allelic diversity, and not strictly by the number of genes at a given isotype; for example, methionine has a low number of encoding tRNAs but a high number with predicted nonfunctional alleles (**Figure 2**).

We observed several instances in which the reference genome allele was a poor representative of the overall population. For example, three genes in *C. briggsae*, one in *C. elegans*, and one in *C. tropicalis* harbored alleles observed only in the reference strain, each of which was sub-optimal relative to other alleles (with an RNA folding score up to 20% lower than that of the best allele). This observation emphasizes the relevance of the population as a whole when defining tRNA repertoires. Relatedly, the occurrence of putatively functional allele(s) throughout the population at a given locus increases confidence that the gene is maintained by selection. We examined cases of rare isodecoders in which the gene set derived from the reference genome included a single instance of a codon in one species that was absent from the other two species. Four such instances occurred, as well as a fifth in which the codon was present in two of the three species (**Table S2**). While we hypothesized that these rare codons might reflect reference strain-specific mutations absent from the broader population, in fact, all strains harbored predicted functional alleles—some of which were fixed in the population—of the same isodecoder class at these loci. We also examined the two SeC tRNAs in *C. briggsae*; most organisms encode a single nuclear SeC (Chan and Lowe 2016). One SeC gene was invariant across the population, as in *C. elegans* and *C. tropicalis*, while the other gene had three observed alleles and exhibited missing genotypes in one third of the strains. The rare isodecoder tRNAs and alleles of the second SeC gene exhibited lower RNA folding scores relative to similar genes, raising questions about their biological function, even as their apparent persistence in the population motivates investigation into their origin and potential maintenance.

Alleles with apparent isotype switches might be presumed deleterious, given their potential to misload amino acids (Giege et al. 1998). Correspondingly, of genes with isotype switch alleles that were not fixed in the population, 70-100% exhibited switch alleles at very low frequency (<0.5%) in the three *Caenorhabditis* species (**Figure 2**, small yellow pie pieces). Alternatively, some isotype switches are tolerated and can persist across taxa (Rogers and Griffiths-Jones 2014; Yona et al. 2013). We also observe many such isotype switch alleles at high frequency and fixed within species, perhaps reflecting ancient switches (**Figure 2**, fully or nearly fully yellow pies). We next investigated these observations further.

### Evidence for both contemporaneous and ancient isotype switching in *Caenorhabditis*

We examined isotype switching in the three species, both to assess polymorphism of isotype switching within extant populations and to identify potential ancestral changes that may have arisen in a common ancestor of one or more *Caenorhabditis* species (∼25 mya). An ‘Isotype switch’ allele may arise by mutation to the anticodon or by alteration of the backbone sequence, either of which may induce a mismatch in the identity of the associated amino acid via even a single nucleotide change (Hipps et al. 1995). When isotype switching was variable in the population, we could confidently infer the derived variant(s) in the majority of cases, such that we continued with this analysis (**Figure S4,** Methods).

We characterized isotype switching in terms of the frequency of isotype switching from each amino acid, as well as the frequency of the isotype switch allele(s) themselves (**Figure 3A**). Genes with isotype switch alleles fell into three major categories: 1) ‘all switch’: all alleles were isotype switches (either one or multiple alleles with the same switch); 2) ‘no switch’: no alleles exhibited predicted isotype mismatches; and 3: ‘some switch, some not’: genes with a mix of switching and non-switching alleles (**Figure 3A**). Isotype switches that were fixed within a population occurred from leucine and threonine in all three species; from alanine, serine, and tryptophan in two of three species; and from arginine, glycine, histidine, and isoleucine in species-specific cases (**Figure 3A, Table S5**). The incidence of isotype switching was not well predicted by the number of genes per isotype. For example, tryptophan had among the fewest genes per isotype but showed fixed or variable isotype switching in all three species, while proline had the near-highest gene count but only exhibited low-frequency variable switching in *C. briggsae* (**Figure 3A**). Per species, the degree of fixed isotype switches, presumably arising deeper in time from a common ancestor, did not correlate with species-wide measures of genetic or tRNA-specific variation: *C. elegans* (intermediate variation) had 7 amino acids with fixed switches, *C. briggsae* (highest variation) had 5, and *C. tropicalis* (lowest variation) had 4 (**Figure 3A**), corresponding to 19, 31, and 25 isotype-switched tRNA genes, respectively.

**Figure 3.**
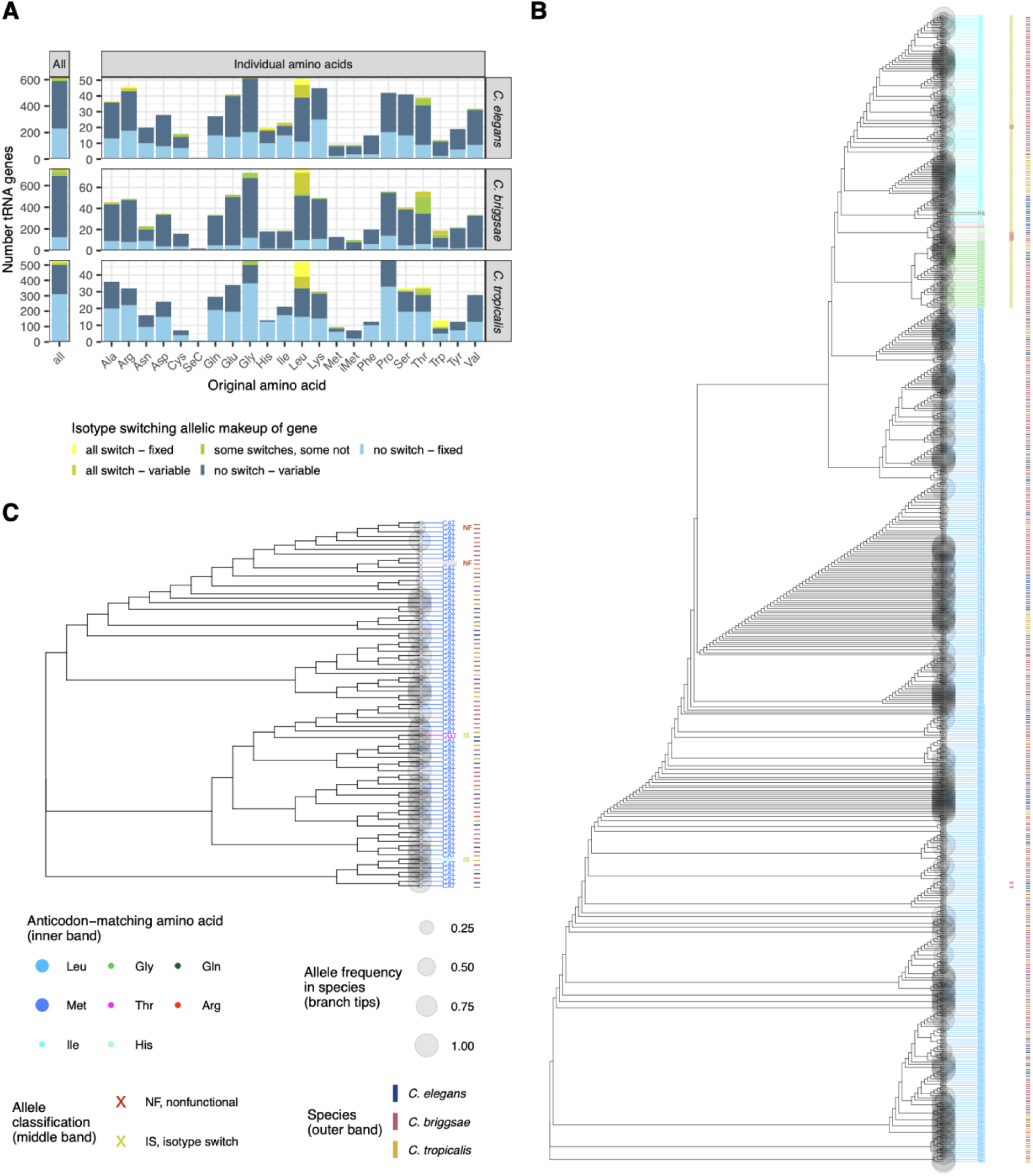
Isotype switching from ancient and contemporaneous tRNA genetic variation. **A,** The number of tRNA genes predicted to encode each amino acid, colored by whether the gene’s anticodon matches the prediction and whether there is allelic variation at that gene in the species. Color definitions: all switch – fixed: a single isotype-switching (anticodon-backbone mismatch) allele is present in all strains; all switch – variable: all alleles harbor isotype switches, but there are multiple alleles in the population; some switches, some not: some alleles observed in the population harbor isotype switches while others do not (variable isotype switching); no switch – variable: no alleles in the population harbor isotype switches, but there are multiple alleles in the population; no switch – fixed: one allele without an isotype switch is fixed in the population. **B** and **C**, neighbor-joining allele cladograms from whole-gene secondary-structure aware tRNA multiple sequence alignments (Methods) for all alleles from all species for genes predicted to have a leucine (**B**) or methionine (not iMet) (**C**) backbone. Branch label color reflects the amino acid encoded by the anticodon, with isotype switches noted in the central circle as yellow and predicted non-functional alleles noted in the central circle as red. In **B**, ancestral leucines are now predicted to [match] isoleucine (cyan) in all strains in related alleles in all three species and glycine (green) in *C. briggsae* and *C. elegans*. In **C**, ancestral methionine now matches threonine (magenta) in a minority of strains in *C. elegans* and isoleucine (cyan) in a minority of strains in *C. briggsae*.

To further evaluate fixed isotype switches, we generated secondary structure-aware multiple sequence alignments of all alleles per amino acid backbone across species (Methods). These alignments enabled visualization of allele sequence relationships. For example, genes with leucine backbones switched to isoleucine in all strains of all three species, and these isoleucine-encoding alleles cluster together across species rather than with leucine-retained alleles from their respective species (**Figure 3B**). Similarly, another set of alleles with leucine backbones showed switches to glycine in all three species and clustered together across species (**Figure 3B**). These examples support the conclusion that some switches arose from ancient, shared events and may be neutral or beneficial. Notably, one gene with a leucine backbone (*C. elegans* I.trna17) exhibited isotype switch alleles for every strain, but 80.2% of the strains carried alleles with a glutamine-recognizing anticodon (TTG in the DNA) while the remaining 19.8% carried alleles with anticodons for isoleucine (TAT).

In genes polymorphic for switching (‘some switch, some not’), the switches tended to be species-specific and low frequency, possibly reflecting relatively recent, deleterious mutations that may be under purifying selection (**Figure 3A, Table S6**). The tRNAs for methionine exemplified this trend (**Figure 3C**). Methionine switched to threonine in a singleton (observed in one strain) allele in *C. elegans* and to isoleucine in a singleton allele in *C. tropicalis* (**Figure 3C**). These rare alleles did not cluster closely on the allele tree, suggesting they arose as independent, relatively recent events (**Figure 3C**). Across species, the extent of low-frequency isotype switching generally mirrored overall genetic and tRNA allelic variation: *C. briggsae* showed the most and *C. tropicalis* showed the least (**Figure 3A**). This observation supports the idea that isotype switching may arise from the high mutation load imposed on tRNAs by TAM; indeed, anticodons frequently harbor genetic variation (**Figure 1C**).

We observed only one gene in which isotype switching was neither rare nor fixed (*C. elegans* tRNA gene V.trna22) (**Table S6**). This backbone sequence of this gene predicted valine and, for 546 strains (79.7% of the population), exhibited an anticodon corresponding to methionine (CAT), including the reference strain. The remaining 118 strains (12.3%) carried alleles without an isotype switch and encoded a valine anticodon (CAC) that matches the backbone. This example highlights the potential for studying isotype switching in *Caenorhabditis* species, from ancient events to contemporaneous variation.

### Distinctive genomic organization of tRNA genes

Genomic context can provide important clues about evolutionary processes. Here, we examined how tRNA genes were organized relative to each other and within the architecture of *Caenorhabditis* genomes. For these analyses, we used the positions of tRNA genes derived from the reference genomes.

We first asked whether tRNA gene distribution followed the same location rules as protein-coding genes in *Caenorhabditis* genomes. The holocentric chromosomes of these species exhibit distinct recombination domains, with higher recombination rates in chromosome arms and lower rates in chromosome centers (Barnes et al. 1995; Hillier et al. 2007; Rockman and Kruglyak 2009; Ross et al. 2011; Kaur and Rockman 2014; Bernstein and Rockman 2016; Noble et al. 2021; Stevens et al. 2022). Commensurately, overall gene density is generally higher and genetic variation lower in centers relative to arms (Barnes et al. 1995; Hillier et al. 2007; Rockman and Kruglyak 2009; Noble et al. 2021), a pattern visible by eye for protein-coding genes (**Figure 4A**). However, tRNA genes did not follow this rule: they were distributed more evenly across chromosomes, sometimes even enriched in chromosome arms (**Figure 4A**). This tRNA-specific gene distribution across recombination domains was evident, and significantly different from that of protein-coding genes, on each of the six chromosomes in each species (**Figure 4A**).

**Figure 4.**
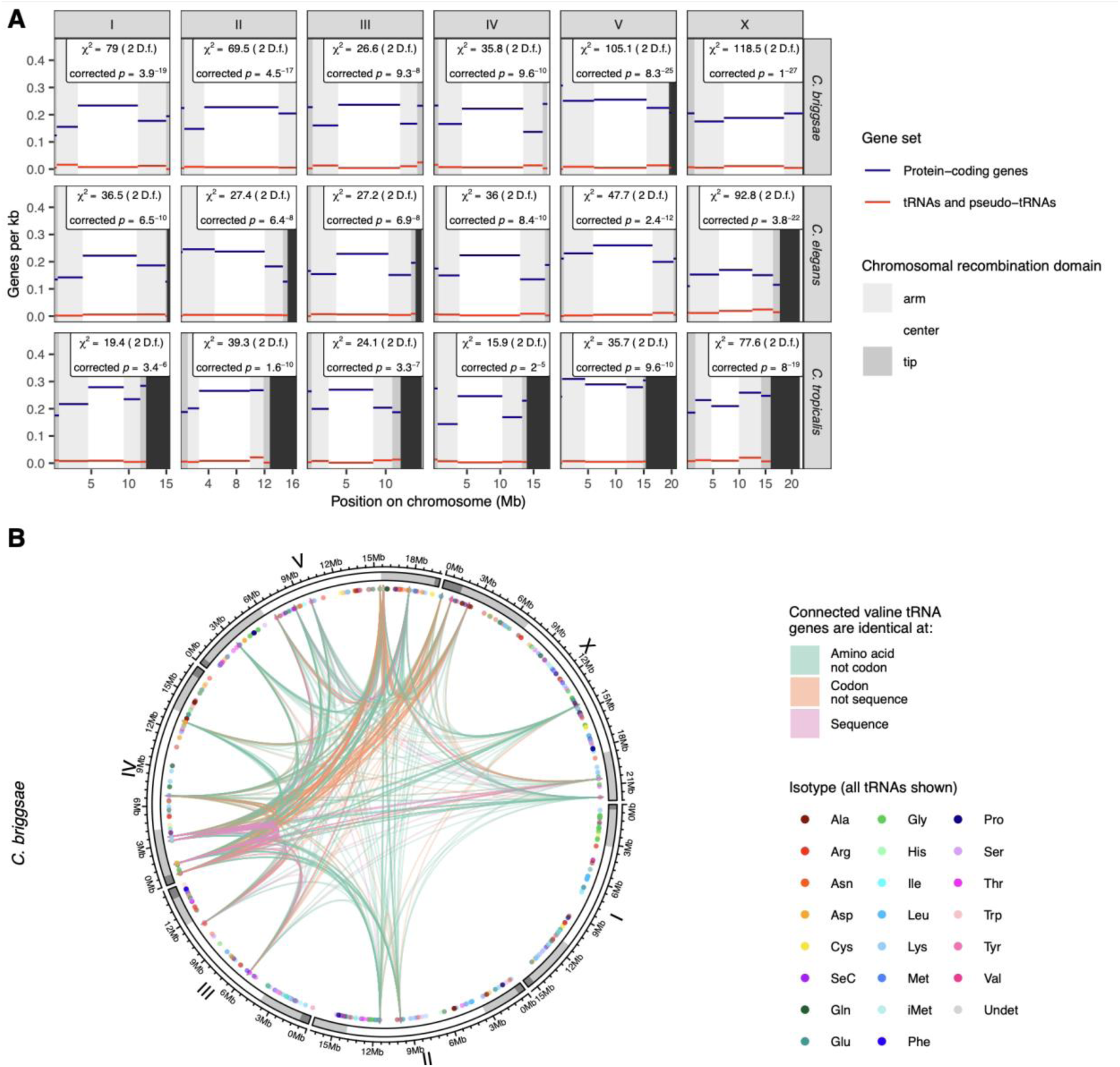
Distribution of tRNA genes within *Caenorhabditis* genomes. **A,** Per-kb density of protein-coding and tRNA genes in each chromosome’s high recombination (arms) and low recombination (center) domains. Species’ chromosomes have different lengths; where one species has a longer chromosome than another, the other species’ plotting region is blacked out. Results from per-species, per-chromosome χ^2^ tests comparing protein-coding and tRNA gene distribution across the domains are annotated on the plot; p-values are Bonferroni corrected for 18 (chromosomes x species) tests. **B,** The distribution of all tRNA genes in the *C. briggsae* reference genome (colored dots, *n* = 779 non-all-nonfunctional allele, single predicted amino acid backbone tRNAs) shown with pairwise connections between all valine tRNA genes (colored lines, *n* connections = 595 between *n* = 35 valine tRNA genes). Chromosomes and genomic coordinates are shown in circular layout; chromosome shading corresponds to chromosomal recombination domain (see **A**).

In all three species, tRNA genes occurred on all chromosomes in all center and arm recombination domains (**Figure 4, Figure S5**). All isotypes comprising more than one tRNA gene had genes on at least half of all chromosomes (three of six) (**Figure 4, Figure S5**). Given this genome-wide distribution, we next asked whether similar tRNA genes were more likely to reside near each other, or if gene locations were random with regard to identity at the isotype, isodecoder, or full sequence level. tRNAs showed evidence of a dispersed, non-stereotyped genomic distribution, as numerous genes with shared identity, including identical sequences, occurred on different chromosomes. For example, in each species, nine genes for initiation methionine (iMet) were found on three (*C. tropicalis*) or four (*C. elegans, C. briggsae*) chromosomes. In *C. elegans* and *C. tropicalis*, all nine iMet tRNA genes had identical sequences in the reference genome; in *C. briggsae*, two distinct sequences corresponded to iMet. Other isotypes, such as valine (**Figure 4B, Figure S5**), contained multiple sequence classes, with both highly similar genes (identical sequences) and more divergent genes (different isodecoders) residing on different chromosomes from one another.

Nevertheless, in all three species, tRNA gene pairs were more likely to reside on the same chromosome the more similar they were: genes within an isotype were more likely to share a chromosome than genes with different isotypes; genes within an isodecoder class (same anticodon) were more likely to share a chromosome than genes within the same isotype but with different anticodons; and genes with full-length identical sequence were more likely to share a chromosome than genes in the same isodecoder class that had other (non-anticodon) sequence differences (**Figure S6**). Furthermore, within the same chromosome, tRNA gene pairs tended to be closer together the more similar they were. This pattern followed the same hierarchy as above, and also showed that sequence similarity correlated with genomic proximity (**Figure S7**).

## Discussion

The universality of tRNAs, including their essentiality, organization in eukaryotic nuclear genomes, and vulnerability to TAM, motivate comprehensive, high-resolution characterization of their allelic variation—in any representative system. *Caenorhabditis* nematodes offer an ideal opportunity, including hundreds of genetically diverse strains and extensive genetics tools developed for the model organism *C. elegans*, which will maximize tractability in follow-on experiments (Barriere and Felix 2005a; Barriere and Felix 2005b; Félix et al. 2013; Crombie et al. 2019; Andersen and Rockman 2022; Crombie et al. 2024).

### Perspective

Our nascent understanding of natural genetic variation in tRNA repertoires, which this work aims to advance, points to unresolved questions that are both broad and biologically significant. For one, does standing variation at tRNA loci confer functional differences among individuals and costs to fitness (Thornlow et al. 2018)? Here, we show that the tRNA pools of three *Caenorhabditis* species harbor extensive intra-specific variation in tRNA genes, including substitutions at every site along the generic molecule, frequency spectra consistent with deleterious effects, and derived mismatches between the amino acid decoded by the anticodon and that specified by other identity elements. In a parallel study, in which we introduced a novel sequence substitution model for TAM, we demonstrated that standing mutations in *C. elegans* cytosolic tRNAs are consistent with the TAM signature and destabilize molecular structure by disproportionately affecting paired sites and reducing thermodynamic stability (Banos et al. 2025). These findings support the hypothesis that tRNA loci contribute meaningful mutational load, though experimental assays will be required to test how these mutations affect molecular and organismal fitness.

Mutations at tRNAs that occur as single-copy genes (e.g., tRNAs for selenocysteine and mt-tRNA) demonstrate strong deleterious effects (Florentz et al. 2003; Yarham et al. 2010; Suzuki et al. 2011; Schoenmakers et al. 2016; Lant et al. 2019), but the high copy number and multidimensional variation of nuclear encoded tRNAs add significant biological and analytical complexity. Work connecting specific molecular effects of tRNA modifications (such as (Torres et al. 2019; Davey-Young et al. 2024; Tennakoon et al. 2025)) to larger-scale organismal phenotypes in experimentally tractable systems across multiple generations will be particularly valuable.

A second question arising from intraspecific variation at tRNA genes relates to their evolution over intermediate timescales. tRNA pools exhibit remarkable evolutionary churn in gene number, sequence, and isotype identity across evolutionary lineages (Rogers et al. 2010; Wang and Ruvinsky 2012; Rogers and Griffiths-Jones 2014; Velandia-Huerto et al. 2016; Yona et al. 2013). These evolutionary patterns and the processes that drive them remain relatively underexplored, in part due to the same obstacles that impede functional analysis of cytosolic tRNAs: the large number of tRNAs in nuclear genomes and their complex organization. Gene orthology is a challenge to resolve syntenically across lineages, due to sequence similarity between loci as well as genuine gene gains and losses (Lant et al. 2019; Velandia-Huerto et al. 2016; Wang and Ruvinsky 2012). However, intraspecific variation in tRNA repertoires offers an entry point to elucidating how these gene sets diverge over time, by revealing early-stage changes that may include gene duplications, losses, or conversions.

Allelic variation at tRNA loci also raises the question of whether some tRNA genes are more critical than others. Recent work in mammals (Torres 2019; Hughes et al. 2023; Gao et al. 2024) and yeast (Bloom-Ackermann et al. 2014) suggests that specific genes within isoacceptor and isodecoder classes exert disproportionate functional impact, despite evidence for functional redundancy (Leung et al. 1991; Kutter et al. 2011). Elucidation of such roles is directly relevant to understanding the fitness consequences of mutational variation, the proximate mechanisms governing tRNA maintenance and decay, and the divergence of tRNA repertoires over time. One approach to this question is to integrate genetic variation data with computational models that consider tRNA gene chromatin conformation, predicted or empirical expression, and canonical tRNA functionality (*e.g.,* (Thornlow et al. 2020)) to predict context-specific tRNA gene function for empirical testing. tRNA gene variants may themselves alter expression, as many tRNAs have internal promoters (Galli et al. 1981; Giege et al. 1998); they are also likely to affect post-transcriptional regulation and activity, including by post-transcriptional modification (Schimmel 2018). Direct quantification and epigenetic profiling, using methods that distinguish identical genes via flanking sequence awareness, in individuals with diverse allelic repertoires and across multiple tissues and life stages, will be informative. Relatedly, mutational variation at tRNA genes can be leveraged to investigate codon usage bias within populations. tRNA allelic combinations may be more or less favored in the context of overall codon usage or within specific gene sets; codon usage itself might also vary and evolve within a species.

Transfer RNAs are uniquely universal. The structure and function of the tRNA molecule are conserved across the tree of life, and in multicellular eukaryotes, nuclear genomes encode hundreds of cytosolic tRNAs (Bermudez-Santana et al. 2010; Goodenbour and Pan 2006). The human genome encodes over 400 putatively functional cytosolic tRNA genes, characterized by complex multidimensionality: genes exhibit both shared and diverged sequence identities and diversity in the number of genes within each isoacceptor class (Goodenbour and Pan 2006; Chan and Lowe 2016). This complexity in the human tRNA pool has been hypothesized to be adaptive, evolving to meet the proteomic demands of humans as highly differentiated multicellular organisms (Pinzaru and Tavazoie 2023). Indeed, more complex organisms, like humans and chimpanzees, harbor more isodecoders than those that are less complex, like flies and worms (Orellana et al. 2022; Goodenbour and Pan 2006). Yet, this modest trend scales poorly with complexity. At only ∼1000 cells (Sulston and Horvitz 1977), *C. elegans* is approximately 30 billion times smaller than a human but encodes over 500 tRNA genes and exhibits similar multidimensionality in distribution of isoacceptors (Duret 2000). While tRNA repertoires do exhibit lineage-specific diversification (McDonald et al. 2025), their general similarities suggest that tRNA functionality and the evolutionary pressures shaping tRNA pools are broadly conserved across eukaryotes and that discoveries in any system will be widely informative.

### Comments on this study

A central phenomenon in tRNA evolution is transcription-associated mutagenesis (TAM), to which tRNA genes are highly exposed (Thornlow et al. 2018). Here, we replicate initial reports in other eukaryotes to show evidence of elevated TAM at tRNA loci in *C. elegans*, *C. tropicalis,* and *C. briggsae* (**Figure 1C**), further emphasizing the universality of this phenomenon (Thornlow et al. 2018). These findings complement our related work, in which we used a novel sequence substitution model to assess mutational variation in *C. elegans*, including its effects on molecular secondary structure and thermostability (Banos et al. 2025). Because tRNA genes are transcribed in all cells, TAM presumably occurs organism-wide, consistent with the high somatic mutation burden in tRNA genes in human cancer and aging (Murillo-Recio et al. 2025). However, only mutations arising in germline cells contribute to the heritable variation observed within and across populations. This reasoning suggests that TAM is universal despite diverse mechanisms of eukaryotic germline formation, reinforcing predictions that population-level variation in tRNA genes imposes a mutation load on genomes across taxa.

The tRNA allelic variation we report here may compromise tRNA function by affecting folding and structure or amino acid proofreading or identity (**Figure 2**). Our analysis indicates that strains differ in the number of functional tRNA genes. However, because we analyzed only tRNA genes present in the reference genome and did not assess copy-number variation at these loci or search for novel tRNAs with distinct sequences or genomic locations in non-reference strains, we cannot explicitly define the number of predicted functional tRNA genes for each strain. Given that copy-number variation of tRNA genes is common, its analysis will be helpful for future work (Iben and Maraia 2012, 2014; Darrow and Chadwick 2014; Velandia-Huerto et al. 2016; Hughes et al. 2023). Complete resolution of cytosolic tRNA complements for individual strains requires strain-specific assemblies, and must distinguish true genomic differences from assembly artifacts that can arise even between assemblies of the same strain or organism (Murillo-Recio et al. 2025).

Isotype switching is a particularly compelling aspect of tRNA variation, considering its potential to compromise the protein sequence. We observed two broad categories of isotype switching (**Figure 3**). First, we detected isotype mismatches that were fixed in the population, which likely represent ancestral changes and contribute to the isotype remolding reported across eukaryotes (Rogers and Griffiths-Jones 2014; Velandia-Huerto et al. 2016). Leucine-backbone tRNAs most often carried switches, with many genes fixed for other anticodons in all three *Caenorhabditis* species. These switches may predate speciation and may have fixed neutrally or adaptively, consistent with observations of leucine tRNAs in nematodes, humans, and other taxa (Lant et al. 2019; Rogers and Griffiths-Jones 2014); confirmation requires resolving orthology and synteny. Because the aminoacyl-tRNA synthetase for leucine, like those for alanine and serine, does not rely on the anticodon for amino acid identity, leucine tRNAs can and do deliver leucine even with anticodons for other amino acids (Giege et al. 1998; Davey-Young et al. 2024). We also observed fixed isotype switches from alanine and serine backbones. Second, we detected polymorphic isotype switches, present in a subset of strains, in all three species. These variants were rare, occurred at low frequency, and based on their tRNA backbones, are unlikely cause widespread amino acid mischarging or misincorporation but could slow amino acid loading and increase energy-intensive proofreading by aminoacyl-tRNA synthetases (Perona and Gruic-Sovulj 2014). Taken together, we predict that the tRNA complements of some strains are suboptimal due to isotype switching; some strains are burdened by each isotype switch, others are not. Determining the functional consequences of these polymorphic switches will clarify the contributions of tRNA gene genetic variation to organismal fitness and mutational load.

The standing variation at tRNA genes that we report here may persist longer in these three nematode species because of their androdioecious mating system; extended selfing with low rates of outcrossing promotes the accumulation of mutational load (Loewe and Cutter 2008). Background selection dominates in these genomes, but varies between the low-recombination centers of chromosomes and the higher-recombination arms (Rockman et al. 2010; Andersen et al. 2012). We show that tRNAs are disproportionately distributed on arms (**Figure 4**), which may reflect an adaptation for more efficient purging of deleterious mutations. This distribution may also facilitate the rapid birth-death cycles that characterize tRNA gene families (Nei and Rooney 2005; Wang and Ruvinsky 2012), given the greater potential for duplication, excision, and conversion under higher crossover rates. tRNAs likely duplicate as mobile genetic elements; indeed, many SINE elements originate from tRNA genes. Different mobile element-derived sequence families have different distributions in the *C. elegans* genome, with some center-biased and others arm-biased (Surzycki and Belknap 2000; Laricchia et al. 2017), so the tRNA-specific distribution might reflect tRNA-specific evolution.

The genetic variation in tRNAs we report here may confer biological significance beyond their canonical role in translation; alternative activities of tRNAs continue to be discovered (reviewed in (Schimmel 2018)). Differences in tRNA genes may affect tRNA participation in protein complexes and the generation of tRNA fragments; variation in human isodecoder gene sets has been observed to impact expression levels of tRNA fragments, which have diverse roles beyond those of full-length tRNAs (Torres 2019; Torres et al. 2019). As previously mentioned, the role of genetic variation on tRNA expression and regulation, including via post-transcriptional modifications, remains an outsized and essential question.

## Conclusion

Here, we uncovered remarkable genetic variation in nuclear-encoded tRNA genes in three *Caenorhabditis* species, much of which may affect tRNA function. The insights, datasets, and resources presented here, including an interactive web app available at https://wildworm.biosci.gatech.edu/trnarefalleles/, will serve as a springboard for investigation of the causes and consequences of nuclear tRNA variation across eukaryotes.

## Materials and Methods

The code written for this study is available at https://github.com/paabylab/trnavcfalleles.

### Genomic data sources

The reference genome for *C. elegans* was version ws283 from WormBase (Sternberg et al. 2024), the same as used for short-read sequencing data alignment and variant calling by the *Caenorhabditis* natural diversity resource (CaeNDR, (Crombie et al. 2024)) in the *C. elegans* 687 sample (strain) VCF used here (20250625 release, https://caendr.org/data/data-release/c-elegans/20250625). For *C. briggsae*, reference genome fasta and 719 sample (strain) VCF were downloaded from CaeNDR (20250626 release, https://caendr.org/data/data-release/c-briggsae/20250626; reference genome build from (Stevens et al. 2022)). Likewise, for *C. tropicalis*, we obtained the reference genome fasta and 622 sample (strain) VCF from CaeNDR (20250627 release, https://caendr.org/data/data-release/c-tropicalis/20250627; reference genome build from (Noble et al. 2021)). Specifically, we used the hard-filtered, isotype-only VCFs in all cases; these VCFs include only wild strains that are not genetically identical and only stringently filtered variants.

### Reference genome tRNA identification

To identify tRNA genes and tRNA-like sequences, we ran tRNAscan-SE 2.0 (v2.0.12) (Chan and Lowe 2019; Chan et al. 2021) on reference genomes for the three *Caenorhabditis* species. We ran tRNAscan-SE using default settings, including full logs, progress reports, and isotype-specific modeling by including options -*-log, --progress, --detail*, and *--isospecific*. The tRNAscan-SE output from the reference genomes thus established the set of tRNA genes (and pseudogenes) in our study; it also provided, for each gene: the codon sequence; the cognate amino acid for this codon; the predicted cognate amino acid based on evolutionary analysis of the tRNA backbone; the Infernal (Nawrocki and Eddy 2013) predicted tRNA folding score; a score of goodness-of-fit to the isotype (amino acid) assigned by the codon (related to the predicted backbone); and a note including if tRNAscan-SE predicted the reference sequence to be a pseudogene. We used these outputs rather than data available on gtRNAdb (Chan and Lowe 2016) in order to use the latest reference genomes available and ensure consistency in our data processing and interpretation across genomes and species. We also sought to maximize inclusivity when establishing the initial gene sets from the reference genomes, given our interest in mutational variation in tRNAs. Reference-sequence genes were predicted to be functional tRNAs when they were not called pseudogenes by tRNAscan-SE, had predicted codons, and had Infernal scores > 20 (usually much higher). Following downstream analyses in which we interrogated all alleles for functional status, we were also able to classify genes according to whether none, some, or all their alleles were predicted to be nonfunctional.

### Determining variation in tRNA flanking regions

We first subset each species’s whole-genome VCF to include only the variants within 1 kb of any tRNA gene or tRNA-like sequence identified above using bcftools (v1.11-19) (Danecek et al. 2021), then recorded the identity of each variant that occurred within 40bp up or downstream of each tRNA gene; we also tracked the strains in which this variant was observed or had missing genotype. Both SNVs and INDELs were included. (Note that all three species are androdiecious and self-mate in nature, so the wild isotypes are naturally inbred homozygous genome-wide; for this reason, we counted any variant called as heterozygous as missing a genotype in that strain.) From these data, we computed the proportion of tRNAs with an observed variant at each flanking basepair from −40 upstream of the gene to 40 downstream of the gene (appropriately accounting for strandedness), overall and restricting to certain gene sets and mutation types.

Code relevant for these analyses is available in our GitHub repository in directory *flank_specific*.

### Allele frequency analyses of variants in tRNAs and protein-coding exon regions

To characterize SNV and INDEL variants in tRNA genes and protein-coding exon regions, we extracted all biallelic variants in these regions from the CaeNDR VCFs. We note that while tRNA genes are highly similar, they are short, so the short-read 90+ bp paired-end Illumina sequencing data underpinning the variant calls will include both tRNA genes and their more unique genomic contexts in the majority of cases. tRNA gene regions were all those in the .bed files generated by tRNAscan-SE (tRNA and tRNA-like genes), and we generated protein-coding exonic regions .bed files from the species’ GTFs. We used bcftools (v1.11-19) (Danecek et al. 2021) to extract the counts of strains with homozygous reference, homozygous alternate, heterozygous (rare in these isogenic species), and missing genotypes for each biallelic variant in a region of interest, retaining the region’s identity. We then used these genotype counts to investigate minor allele frequency (MAF) distributions between tRNA and protein-coding regions; these comparisons’ results were robust to the calculation of MAF as the number of minor allele-homozygous genotypes divided by the total number of strains or divided by the number of strains with non-missing genotypes at that particular variant. Tajima’s *D* (Tajima 1989) was calculated from the site frequency spectrum generated from these genotype counts (following (Yang 2022)). We performed this calculation restricting for multiple site subsets as missingness impacts the calculation: sites with >80% or >90% samples with non-missing genotypes. We also performed the calculation based only on SNVs or also including INDELs; results were robust to all variations.

Code relevant for these analyses is available in our GitHub repository in the directory *variant_characterization*.

### Gene location analyses

Analyses of gene location with regard to recombination domain used recombination domain genomic boundaries for *C. elegans* from (Rockman and Kruglyak 2009) (Table 1), for *C. briggsae* from (Stevens et al. 2022) (Table 2), and for *C. tropicalis* from (Noble et al. 2021) (‘ctbp’ data in https://github.com/lukemn/tropicalis/blob/master/geneticMap/data/c_tropicalis_flye_geneticMap.rda). All protein-coding genes for each species for comparison were obtained from the GTF files matching the reference genomes (ws283 from WormBase, narrowed to the protein-coding biotype for *C. elegans*; protein-coding only gene models generated by CaeNDR for *C. briggsae* and *C. tropicalis,* canonical_geneset.gtf files). We assigned genes to specific recombination domains based on the domain that encompassed the midpoint of the gene. From these assignments, we computed the per-kb gene density of each class of gene in each recombination region; the difference in domain distribution between genes within species and chromosomes (*i.e*., arm vs center gene density); and the difference in domain distribution between the gene sets within species and chromosomes (*i.e*., differential arm vs center gene density between two species).

To examine the pairwise location distribution of tRNA genes, we used the gene’s midpoint as its single location value and, for each pair of tRNA genes in the reference genome, determined whether the pair was on the same chromosome and, if so, how far apart the midpoints were.

Then each pair was classified in various ways relevant to its analysis (whether the gene pair had identical sequence; whether the gene pair was from the same isotype; the sequence alignment divergence between the gene pair [see below]; etc).

Code relevant for these analyses is available in our GitHub repository: *analysis/trna_location_analyses.R*.

### tRNA allele identification and characterization

To identify all tRNA alleles in the three populations, we narrowed the VCFs to include only tRNA-like sequences and surrounding regions (identified by tRNAscan-SE as described above), then generated allele-specific FASTA files by inserting each observed variant(s) for each strain, overwriting the reference base at that position (SNVs and INDELs included). Strains with missing genotypes at any variant within a tRNA were not assigned an allele identity for that gene and were instead recorded as missing. To characterize the alleles, we ran tRNAscan-SE on the FASTA file containing all tRNA allele sequences (options as used for the reference genomes). Therefore, each allele was annotated with its predicted isotype (codon-based) and amino acid backbone, as well as all other tRNAscan-SE output information. Each allele was given predicted functional classifications based on the following criteria:

1. ‘Best’: an allele was classified as ‘best’ if its isotype matched its predicted amino acid backbone (identical ‘AA’ and ‘AlleleCM’ columns in tRNAscan-SE output) and it had the best (highest and putatively functional) infernal score of any allele at this tRNA (and was not called a pseudogene by tRNAscan-SE or missed in tRNAscan-SE’s calling)
2. ‘Functional’: as ‘Best’, but with a lower infernal score (still >>20)
3. ‘Isotype switch’: an allele was classified as ‘isotype switch’ if it was presumed functional (not meeting any ‘nonfunctional’ criteria, below) and its isotype did not match its predicted amino acid backbone (‘AA’ and ‘AlleleCM’ columns in tRNAscan-SE output differed)
4. ‘Nonfunctional’: an allele was classified as ‘nonfunctional’ if it was called a pseudogene by tRNAscan-SE, missed in tRNAscan-SE’s calling, had an indecipherable/NNN codon, or had an infernal score < 20 (in practice, all with such scores were flagged as pseudogenes).

These calls were used in further gene-level analyses, such as identifying genes with all pseudogenized/nonfunctional alleles and classifying genes as having fixed isotype switching or variable isotype switching.

To estimate confidence in our isotype switching classifications specifically, we intersected observed genetic variation (from the VCF) with tRNAscan-SE-derived allele annotations. At genes where all alleles were called isotype switches, no genotype variability exists, so we could not interpret how often we could pinpoint genotype changes ‘used’ by tRNAscan-SE. Therefore, we focused on genes with segregating alleles that differ in their cognate amino acid/isotype switch identity. Among these, 56-61% of variable (low-frequency) isotype switches were attributed to anticodon mutations and 39-44% to backbone mutations. In all backbone mutation cases and 58-80% of the anticodon isotype switch cases, we could clearly map variants in the VCF to the predicted isotype switch. Some of the unexplained isotype switch calls likely stem from insertion-deletion polymorphisms, which we included in tRNA allele generation but excluded from variant analysis. Combined with the fact that most high-frequency isotype switches were anticodon-based and necessarily excluded from this analysis, we infer that these numbers represent a conservative lower bound on the reliability of isotype switch identification.

Code relevant for these analyses is available in our GitHub repository: see especially all scripts in *generate_alleles*; *analysis/show_mutational_variation.R; analysis/charisotypeswitching.R*.

### Secondary-structure aware tRNA sequence alignment and related analyses

To enable secondary structure-aware alignment of tRNA gene sequence, as these structures do not have invariant lengths across sequences (Chan and Lowe 2016), we used the secondary structure predictions from tRNAscan-SE to determine for each tRNA gene in the reference genome and each allele which base mapped to which structure. We first excluded introns from this analysis, excising the tRNAscan-SE predicted intron positions from the tRNA gene sequence. For example, we first identified the codon, as tRNAscan-SE gives its position in its secondary structure output, then we inferred the surrounding unpaired regions were part of the anticodon loop, the paired regions surrounding them were part of the anticodon stem region, etc. In some cases, especially pseudogenes, we could not map all bases in the tRNA to a predicted secondary structure, so we flagged these and excluded them from downstream analyses. In other cases, predictions were imperfect, for example, not all bases in the acceptor stem were paired; since this can and does happen in biology, we retained these in analyses while flagging them. We determined that including vs. excluding all genes with such tags did not yield obviously different results, so we included them in the results provided here.

We used the same logic to identify the relative position within secondary structure of each VCF variant: we broke the reference tRNA up into its secondary structure components, mapped genomic positions to the position of each component, and recorded the relative position in the relevant secondary structure. This enabled us to generate estimates of the per-structure proportion of tRNAs with variants and normalize this to the average length of the structure across all tRNAs.

We generated multiple sequence alignments of all alleles observed across all three species populations in a cloverleaf-structure aware fashion by breaking each sequence into its individual structural components as specified above, then aligning each structure separately, and finally concatenating the per-structure alignments into full tRNA alignments. Specifically, we ran *foursale* (4SALE CMD version, v1.3) (Wolf et al. 2014), which uses ClustalW2 (v2.1) (Larkin et al. 2007) to align sequences after combining the RNA secondary structure information and base information at each base. Parameters were -*gapextend 1* and *-gapopen 1*. This was run separately for each consecutive region of the tRNA: the acceptor stem’s left (first) paired part to the beginning of the D arm; the D arm (stem and loop) to the beginning of the anticodon arm; the anticodon arm with the anticodon N-masked to avoid anticodon misalignment; separately the anticodon alone (to force anticodon alignment); the variable arm; the T arm up to the acceptor stem; and then the remaining part of the acceptor stem and the acceptor stem overhang. These separate alignments were concatenated in this order, with the anticodon-specific alignment replacing the N-masked anticodon in the anticodon arm alignment, to get the full-length, all-allele tRNA multiple sequence alignment.

These alignments were used 1) for determining the sequence alignment distance for all pairs of unique sequences and 2) for generating allele trees. The sequence alignment distance or sequence divergence was calculated in two ways, only for unique sequences of alleles in the reference genome: first, simply the number of character differences between two alignments (so a gap is counted as different from any base, and different bases are counted as different from each other); second, the number of distinct stretches of difference between two alignments. Neighbor-joining trees were generated from all alleles or subsets of alleles (from within a species; from a specific amino acid background; etc) using maximum-likelihood sequence alignment distances (functions *dist.ml* and *NJ* from the *phangorn* R package (v2.12.1) (Schliep 2011). Trees were further annotated, manipulated, and plotted using the *ape* (v5.8-1) (Paradis and Schliep 2019), *ggtree* (v3.16.3) (Yu et al. 2017), and *tidytree* (v0.4.6)(Yu 2022) R packages.

### General software tools used for analyses and figures

Tools used for specific analytical purposes are often described in the relevant sections; here, we share tools used for general thinking, data processing, and figure creation.

We used Georgia Tech’s Enterprise access to Microsoft Copilot (running GPT 4 and 5) for occasional problem-solving and text editing suggestions (not for any substantial idea or text generation).

Analysis scripts were written in R (up to v4.5.1) (R Core Team 2025) and Python (v3.7) (www.python.org). Workflow scripts were written and run using Nextflow (v22.10.7) (www.nextflow.io). Compute-intensive analyses and workflows were run via the Partnership for an Advanced Computing Environment (PACE), the high-performance computing environment at the Georgia Institute of Technology.

Data wrangling included R packages *data.table* (v1.17.8) (Dowle and Srinivasan 2022), *argparser* (v0.7.2) (Shih 2021), *formattable* (v0.2.1) (Ren and Russell 2021), and *RRNA* (v1.2) (Bida and Maher 2012) and Python packages included pyvcf (v0.6.4) and numpy (v1.23.0) (Harris et al. 2020). Data display and figure creation R packages included *ggplot2* (v3.5.2) (Wickham 2016), *cowplot* (v1.2.0) (Wilke 2020), *RColorBrewer* (v1.1-3) (Neuwirth 2022), *ggforce* (v0.4.0) (Pedersen 2022), *ggsignif* (v0.6.4) (Ahlmann-Eltze and Patil 2021), *ggpattern* (v1.1.4) (FC et al. 2022), *ggpmisc* (v0.6.2) (Aphalo 2024), *ggnewscale* (0.4.2) (Campitelli 2025), *ggh4x* (v0.3.1) (van den Brand 2025), *gridBase* (v0.4.7) (Murrell 2014), *scales* (v1.4.0) (Wickham et al. 2025), *scatterpie* (v0.2.5) (Yu 2025), *circlize* (v0.4.16) (Gu et al. 2014), *magick* (v2.8.7) (Ooms 2025a), and *pdftools* (v3.5.0) (Ooms 2025b).

## Data Availability

Datasets generated in this study are available as supplemental material or via the Zenodo repository at https://doi.org/10.5281/zenodo.17237304. Data is interactively available via web app at https://wildworm.biosci.gatech.edu/trnarefalleles/. Code used in this study is available at https://github.com/paabylab/trnavcfalleles.

## Acknowledgments

The authors thank members of the Southeast Center for Mathematics and Biology, Abigail Lind and members of the Lind lab, Rohini Janivara, and Abbe LaBella for helpful conversations about this work. The Georgia Tech College of Sciences’ Academic and Research Computing Services team provided web server configuration support for the interactive web app. Research cyberinfrastructure resources and services provided by the Partnership for an Advanced Computing Environment at Georgia Tech supported this research. This work was made possible by publicly available data from CaeNDR (https://caendr.org/) and the many members of the research community who have collected and shared strains.

