## Supplemental Tables for "Allelic variation at tRNA genes in three nematode species indicates mutation load despite strong purifying selection"

**Table S1. Number of tRNAs encoding each amino acid in three *Caenorhabditis* species.** Each number is the number of genes of that predicted backbone (tRNAScan-SE AlleleCM) in the reference genome, excluding genes where the reference allele was predicted to be nonfunctional.

| | **Number of tRNA genes** | | |
|---|---|---|---|
| **Amino acid** | *C. elegans* | *C. briggsae* | *C. tropicalis* |
| Ala | 37 | 48 | 36 |
| Arg | 45 | 48 | 36 |
| Asn | 20 | 22 | 16 |
| Asp | 28 | 35 | 24 |
| Cys | 14 | 16 | 10 |
| SeC | 1 | 2 | 1 |
| Gln | 27 | 36 | 27 |
| Glu | 41 | 54 | 38 |
| Gly | 51 | 80 | 50 |
| His | 20 | 18 | 14 |
| Ile | 24 | 29 | 23 |
| Leu | 50 | 78 | 54 |
| Lys | 46 | 54 | 37 |
| Met | 9 | 13 | 11 |
| iMet | 9 | 10 | 9 |
| Phe | 15 | 21 | 14 |
| Pro | 42 | 57 | 51 |
| Ser | 40 | 41 | 31 |
| Thr | 33 | 40 | 32 |
| Trp | 12 | 16 | 13 |
| Tyr | 19 | 21 | 12 |
| Val | 32 | 35 | 31 |

**Tabel S2. Number of tRNA genes with each codon (DNA) in three *Caenorhabditis* species.** Codons are sorted by the amino acid each codon matches (second column; canonical amino acid this codon matches ignoring possible isotype switching). The number is for all genes in the reference genome, excluding genes where the reference allele predicted to be nonfunctional. All theoretical DNA codons are shown regardless of if they appear in any tRNAs in any species.

| Codon (DNA sequence in tRNA gene) | Amino acid | Number of tRNA genes | | |
|---|---|---|---|---|
| | | *C. elegans* | *C. briggsae* | *C. tropicalis* |
| AGC | Ala | 23 | 28 | 20 |
| CGC | Ala | 4 | 7 | 5 |
| GGC | Ala | 0 | 0 | 0 |
| TGC | Ala | 9 | 12 | 11 |
| ACG | Arg | 19 | 23 | 18 |
| CCG | Arg | 2 | 1 | 1 |
| CCT | Arg | 4 | 2 | 3 |
| GCG | Arg | 0 | 0 | 0 |
| TCG | Arg | 10 | 10 | 7 |
| TCT | Arg | 8 | 11 | 9 |
| ATT | Asn | 0 | 0 | 0 |
| GTT | Asn | 20 | 22 | 16 |
| ATC | Asp | 0 | 1 | 0 |
| GTC | Asp | 28 | 35 | 24 |
| ACA | Cys | 1 | 0 | 0 |
| GCA | Cys | 13 | 16 | 10 |
| CTA | SeC | 0 | 0 | 0 |
| TCA | SeC | 1 | 2 | 1 |
| TTA | SeC | 0 | 0 | 0 |
| CTG | Gln | 7 | 11 | 7 |
| TTG | Gln | 21 | 25 | 20 |
| CTC | Glu | 24 | 35 | 24 |
| TTC | Glu | 17 | 19 | 15 |
| ACC | Gly | 0 | 0 | 0 |
| CCC | Gly | 3 | 6 | 4 |
| GCC | Gly | 15 | 20 | 13 |
| TCC | Gly | 36 | 60 | 37 |
| ATG | His | 0 | 0 | 0 |
| GTG | His | 19 | 18 | 14 |
| AAT | Ile | 21 | 24 | 20 |

| | | | | |
|---|---|---|---|---|
| GAT | Ile | 0 | 0 | 0 |
| TAT | Ile | 7 | 25 | 15 |
| AAG | Leu | 20 | 25 | 18 |
| CAA | Leu | 7 | 12 | 7 |
| CAG | Leu | 6 | 7 | 5 |
| GAG | Leu | 0 | 0 | 0 |
| TAA | Leu | 3 | 6 | 5 |
| TAG | Leu | 3 | 7 | 4 |
| CTT | Lys | 34 | 37 | 25 |
| TTT | Lys | 15 | 17 | 12 |
| CAT | Met | 10 | 13 | 11 |
| CAT | iMet | 9 | 10 | 9 |
| AAA | Phe | 0 | 0 | 1 |
| GAA | Phe | 15 | 21 | 14 |
| AGG | Pro | 7 | 6 | 8 |
| CGG | Pro | 4 | 4 | 5 |
| GGG | Pro | 0 | 1 | 1 |
| TGG | Pro | 32 | 47 | 38 |
| ACT | Ser | 0 | 0 | 1 |
| AGA | Ser | 15 | 16 | 12 |
| CGA | Ser | 6 | 8 | 5 |
| GCT | Ser | 10 | 9 | 6 |
| GGA | Ser | 0 | 0 | 0 |
| TGA | Ser | 9 | 7 | 7 |
| AGT | Thr | 17 | 20 | 16 |
| CGT | Thr | 7 | 7 | 5 |
| GGT | Thr | 0 | 0 | 0 |
| TGT | Thr | 12 | 11 | 8 |
| CCA | Trp | 12 | 14 | 9 |
| ATA | Tyr | 0 | 0 | 0 |
| GTA | Tyr | 19 | 21 | 12 |
| AAC | Val | 20 | 24 | 18 |
| CAC | Val | 6 | 6 | 7 |
| GAC | Val | 0 | 0 | 0 |
| TAC | Val | 5 | 5 | 7 |

**Table S3. Metrics relating to the allele frequency spectra of genetic variants in all protein-coding exons, putatively functional tRNA genes, and putatively nonfunctional tRNA genes** (the latter two groups comprise all sequences identified by tRNAScan-SE; see Methods for their definitions)**.** All statistical test p-values shown were corrected for performing two pairwise tests in each case. Abbreviations used in table: MAF, minor allele frequency; SNV, single nucleotide variant; INDEL, insertion-deletion polymorphism.

| | | *C. elegans* | | | *C. briggsae* | | | *C. tropicalis* | | |
|---|---|---|---|---|---|---|---|---|---|---|
| | | Protein coding exons | tRNAs | Non-functional tRNAs | Protein coding exons | tRNAs | Non-functional tRNAs | Protein coding exons | tRNAs | Non-functional tRNAs |
| *Singleton proportion related* | Number variants (SNVs and INDELs) | 859587 | 1360 | 385 | 829946 | 3262 | 618 | 626686 | 794 | 146 |
| | Number singletons (% singletons) | 246591 (28.7%) | 683 (50.2%) | 106 (27.5%) | 307917 (37.1%) | 1738 (53.3%) | 213 (34.5%) | 148307 (23.7%) | 388 (48.9%) | 48 (32.9%) |
| | $\chi^2$ test results, protein-coding exons vs. tRNAs (functional only) | $\chi^2$ = 855516 , $df$ = 1<br>$p$ = 1 x $10^{-92889}$ | | | $\chi^2$ = 820211, $df$ = 1,<br>$p$ = 1 x $10^{-89056}$ | | | $\chi^2$ = 624308, $df$ = 1<br>$p$ = 1 x $10^{-67786}$ | | |
| | $\chi^2$ test results, protein-coding exons vs. functional and nonfunctional tRNAs | $\chi^2$ = 854366 , $df$ = 1<br>$p$ = 1 x $10^{-92764}$ | | | $\chi^2$ = 818378, $df$ = 1<br>$p$ = 1 x $10^{-88857}$ | | | $\chi^2$ = 623871, $df$ = 1<br>$p$ = 1 x $10^{-67739}$ | | |
| *Distribution central tendency related* | Median MAF | 0.0073 | 0.0015 | 0.0073 | 0.0042 | 0.0014 | 0.0042 | 0.0225 | 0.0032 | 0.0064 |
| | Mann-Whitney test results, protein-coding exons vs. tRNAs (functional only) | $W$ = 404758843.5<br>$p$ = 3 x $10^{-88}$ | | | $W$ = 1048948387.5<br>$p$ = 9 x $10^{-116}$ | | | $W$ = 141744761<br>$p$ = 2 x $10^{-99}$ | | |
| | Mann-Whitney test results, protein-coding exons vs. functional and nonfunctional tRNAs | W = 563015113<br>p = 1 x $10^{-74}$ | | | W = 1308498571.5<br>p = 1 x $10^{-95}$ | | | $W$ = 17985348<br>$p$ = 2 x $10^{-96}$ | | |

**Table S4. Tajima’s D in protein-coding exons, putatively functional tRNAs, and putatively nonfunctional tRNAs** (the latter two groups comprise all sequences identified by tRNAScan-SE; see Methods for their definitions)**.** Tajima’s *D* was calculated with several input sets of variants, as noted in the table. Abbreviations used in table: INDEL, insertion-deletion polymorphism

| | | *C. elegans* | | | *C. briggsae* | | | *C. tropicalis* | | |
|---|---|---|---|---|---|---|---|---|---|---|
| Variant inclusion criteria | Metric | Protein coding exons | tRNAs | Non-functional tRNAs | Protein coding exons | tRNAs | Non-functional tRNAs | Protein coding exons | tRNAs | Non-functional tRNAs |
| >=90% strains have genotype, biallelic sites including INDELs | N variants included | 766486 | 1327 | 346 | 753207 | 3106 | 542 | 583407 | 784 | 130 |
| | Tajima's *D* | -1.10 | -2.15 | -1.41 | -1.49 | -2.32 | -1.56 | -0.41 | -2.24 | -1.03 |
| >=90% strains have genotype, biallelic sites excluding INDELs | N variants included | 707446 | 1096 | 294 | 693696 | 2587 | 480 | 545587 | 698 | 106 |
| | Tajima's *D* | -1.07 | -2.12 | -1.35 | -1.45 | -2.30 | -1.48 | -0.37 | -2.21 | -1.02 |
| >=80% strains have genotype, biallelic sites including INDELs | N variants included | 807038 | 1347 | 367 | 787664 | 3182 | 590 | 605533 | 788 | 133 |
| | Tajima's *D* | -1.08 | -2.14 | -1.44 | -1.48 | -2.32 | -1.52 | -0.38 | -2.24 | -1.03 |
| >=80% strains have genotype, biallelic sites excluding INDELs | N variants included | 743117 | 1112 | 308 | 724103 | 2651 | 521 | 565449 | 699 | 108 |
| | Tajima's *D* | -1.04 | -2.11 | -1.37 | -1.44 | -2.30 | -1.43 | -0.35 | -2.21 | -1.03 |

**Table S5. All examples of putative isotype switching observed in this study where isotype switching is fixed in the species at this gene.** Each gene with an isotype switching allele with RNA folding (Infernal) score > 25 is shown. Full per-allele information is available with the source data provided with this study.

| Species | tRNA gene name | tRNA backbone (predicted by tRNAScan-SE) | Amino acid based on codon | Codon | N alleles at this gene with isotype switch | Total number alleles at this gene | RNA folding score of isotype switch alleles (Infernal, median) | N strains with isotype switch allele(s) | Switch allele freq. (of strains with non-missing genotypes here) | Switch allele freq. (of all strains) |
|---|---|---|---|---|---|---|---|---|---|---|
| *C. elegans* | II.trna16 | Ala | Thr | AGT | 1 | 1 | 74.8 | 684 | 100% | 100% |
| | IV.trna13 | Arg | Lys | CTT | 3 | 3 | 26.1 | 194 | 94.63% | 28.36% |
| | IV.trna74 | Arg | Lys | CTT | 1 | 1 | 26.1 | 684 | 100% | 100% |
| | X.trna12 | Arg | Thr | TGT | 3 | 3 | 37.7 | 658 | 100% | 96.20% |
| | X.trna292 | His | Thr | TGT | 1 | 1 | 48.1 | 684 | 100% | 100% |
| | X.trna259 | Ile | Lys | CTT | 2 | 2 | 27.6 | 8 | 1.17% | 1.17% |
| | I.trna10 | Leu | Ile | TAT | 1 | 1 | 84.7 | 684 | 100% | 100% |
| | I.trna18 | Leu | Ile | TAT | 3 | 3 | 76.9 | 675 | 100% | 98.68% |
| | I.trna23 | Leu | Gly | CCC | 1 | 1 | 79.1 | 684 | 100% | 100% |
| | I.trna54 | Leu | Gly | CCC | 2 | 2 | 69.25 | 635 | 100% | 92.84% |
| | I.trna55 | Leu | Gly | CCC | 2 | 2 | 79.15 | 681 | 100% | 99.56% |
| | I.trna57 | Leu | Ile | TAT | 1 | 1 | 84.7 | 684 | 100% | 100% |
| | III.trna60 | Leu | Arg | CCT | 1 | 1 | 68.7 | 684 | 100% | 100% |
| | III.trna61 | Leu | Lys | CTT | 3 | 3 | 70.2 | 679 | 100% | 99.27% |
| | IV.trna25 | Leu | His | GTG | 2 | 2 | 78.7 | 684 | 100% | 100% |
| | X.trna227 | Leu | Ile | TAT | 4 | 4 | 77.85 | 682 | 100% | 99.71% |
| *C. briggsae* | I.trna27 | Ala | Leu | TAG | 5 | 5 | 49.3 | 502 | 100% | 69.82% |
| | I.trna116 | Leu | Ile | TAT | 6 | 6 | 76.9 | 704 | 100% | 97.91% |
| | I.trna119 | Leu | Ile | TAT | 6 | 6 | 71.5 | 712 | 100% | 99.03% |
| | I.trna120 | Leu | Ile | TAT | 6 | 6 | 78.1 | 696 | 100% | 96.80% |
| | I.trna121 | Leu | Ile | TAT | 2 | 2 | 75.55 | 710 | 100% | 98.75% |

| | | | | | | | | | | |
|---|---|---|---|---|---|---|---|---|---|---|
| | I.trna122 | Leu | Ile | TAT | 4 | 4 | 76.05 | 714 | 100% | 99.30% |
| | I.trna123 | Leu | Ile | TAT | 4 | 4 | 74.65 | 667 | 100% | 92.77% |
| | I.trna124 | Leu | Ile | TAT | 1 | 1 | 79.4 | 719 | 100% | 100% |
| | I.trna28 | Leu | Ile | TAT | 2 | 2 | 65.7 | 719 | 100% | 100% |
| | I.trna29 | Leu | Ile | TAT | 4 | 4 | 70.65 | 669 | 100% | 93.05% |
| | I.trna30 | Leu | Ile | TAT | 4 | 4 | 76.85 | 711 | 100% | 98.89% |
| | I.trna31 | Leu | Ile | TAT | 3 | 3 | 69.5 | 687 | 100% | 95.55% |
| | I.trna32 | Leu | Ile | TAT | 1 | 1 | 59.4 | 719 | 100% | 100% |
| | I.trna33 | Leu | Ile | TAT | 2 | 2 | 64.35 | 714 | 100% | 99.30% |
| | I.trna34 | Leu | Ile | TAT | 1 | 1 | 80.2 | 719 | 100% | 100% |
| | I.trna35 | Leu | Ile | TAT | 8 | 8 | 73.65 | 675 | 100% | 93.88% |
| | I.trna36 | Leu | Ile | TAT | 1 | 1 | 82.7 | 719 | 100% | 100% |
| | I.trna37 | Leu | Ile | TAT | 3 | 3 | 80.2 | 710 | 100% | 98.75% |
| | I.trna38 | Leu | Ile | TAT | 4 | 4 | 66.7 | 719 | 100% | 100% |
| | I.trna39 | Leu | Ile | TAT | 3 | 3 | 65.2 | 715 | 100% | 99.44% |
| | II.trna104 | Leu | Gly | CCC | 4 | 4 | 78.15 | 599 | 100% | 83.31% |
| | II.trna17 | Leu | Gly | CCC | 2 | 2 | 80.35 | 717 | 100% | 99.72% |
| | II.trna18 | Leu | Gly | CCC | 3 | 3 | 74 | 714 | 100% | 99.30% |
| | II.trna19 | Leu | Gly | CCC | 7 | 7 | 76.6 | 688 | 100% | 95.69% |
| | III.trna21 | Ser | Leu | CAA | 7 | 7 | 55.5 | 709 | 100% | 98.61% |
| *C. tropicalis* | I.trna15 | Leu | Ile | TAT | 2 | 2 | 76.4 | 620 | 100% | 99.68% |
| | I.trna16 | Leu | Ile | TAT | 1 | 1 | 78.5 | 622 | 100% | 100% |
| | I.trna17 | Leu | Ile | TAT | 3 | 3 | 77.4 | 619 | 100% | 99.52% |
| | I.trna20 | Leu | Ile | TAT | 1 | 1 | 78.5 | 622 | 100% | 100% |
| | I.trna21 | Leu | Ile | TAT | 2 | 2 | 80.7 | 622 | 100% | 100% |
| | I.trna68 | Leu | Ile | TAT | 1 | 1 | 75.8 | 622 | 100% | 100% |
| | I.trna70 | Leu | Ile | TAT | 1 | 1 | 78.5 | 622 | 100% | 100% |
| | I.trna71 | Leu | Ile | TAT | 3 | 3 | 76 | 620 | 100% | 99.68% |
| | I.trna72 | Leu | Ile | TAT | 1 | 1 | 78.5 | 622 | 100% | 100% |
| | I.trna73 | Leu | Ile | TAT | 2 | 2 | 74.3 | 620 | 100% | 99.68% |
| | I.trna74 | Leu | Ile | TAT | 1 | 1 | 78.5 | 622 | 100% | 100% |
| | I.trna77 | Leu | Ile | TAT | 2 | 2 | 76.45 | 619 | 100% | 99.52% |
| | II.trna88 | Leu | Gly | CCC | 1 | 1 | 75.4 | 622 | 100% | 100% |
| | II.trna89 | Leu | Gly | CCC | 1 | 1 | 69.9 | 622 | 100% | 100% |

| | | | | | | | | | | |
|---|---|---|---|---|---|---|---|---|---|---|
| | II.trna90 | Leu | Gly | CCC | 2 | 2 | 66.7 | 616 | 100% | 99.04% |
| | II.trna91 | Leu | Gly | CCC | 1 | 1 | 69.9 | 622 | 100% | 100% |
| | I.trna22 | Ser | Phe | AAA | 1 | 1 | 27.9 | 622 | 100% | 100% |
| | I.trna4 | Thr | Val | CAC | 1 | 1 | 52 | 622 | 100% | 100% |
| | I.trna66 | Thr | Pro | GGG | 2 | 2 | 44.4 | 621 | 100% | 99.84% |
| | I.trna92 | Thr | Leu | TAA | 3 | 3 | 53.7 | 617 | 100% | 99.20% |
| | III.trna12 | Trp | Arg | CCT | 1 | 1 | 51.5 | 622 | 100% | 100% |
| | III.trna54 | Trp | Gly | GCC | 1 | 1 | 52.7 | 622 | 100% | 100% |
| | III.trna55 | Trp | Arg | CCT | 1 | 1 | 62 | 622 | 100% | 100% |
| | IV.trna13 | Trp | Glu | CTC | 1 | 1 | 63.5 | 622 | 100% | 100% |

**Table S6. All examples of putative isotype switching observed in this study where isotype switching is polymorphic in the species at this gene.** Each gene with an isotype switching allele with RNA folding (Infernal) score > 25 is shown. Full per-allele information is available with the source data provided with this study.

| Species | tRNA gene name | tRNA backbone (predicted by tRNAScan-SE) | Amino acid based on codon | Codon | N alleles at this gene with isotype switch | Total number alleles at this gene | RNA folding score of isotype switch alleles (Infernal, median) | N strains with isotype switch allele(s) | Switch allele freq. (of strains with non-missing genotypes here) | Switch allele freq. (of all strains) |
|---|---|---|---|---|---|---|---|---|---|---|
| *C. elegans* | X.trna140 | Glu | Lys | CTT | 1 | 3 | 70.4 | 2 | 0.29% | 0.29% |
| | X.trna23 | Met | Thr | CGT | 1 | 4 | 77 | 1 | 0.15% | 0.15% |
| | V.trna12 | Thr | Lys | TTT | 2 | 5 | 72.35 | 2 | 0.29% | 0.29% |
| | X.trna13 | Thr | Lys, Arg | CTT, CCT | 2 | 4 | 42 | 323 | 67.71% | 47.22% |
| | X.trna19 | Thr | Ile | AAT | 1 | 4 | 78.8 | 5 | 0.75% | 0.73% |
| | X.trna199 | Thr | Ile | AAT | 1 | 3 | 76.9 | 1 | 0.15% | 0.15% |
| | X.trna299 | Thr | Ile | AAT | 1 | 3 | 85 | 1 | 0.15% | 0.15% |
| | IV.trna33 | Trp | Sup | TCA | 1 | 4 | 71.5 | 1 | 0.15% | 0.15% |
| | V.trna22 | Val | Met | CAT | 2 | 6 | 40 | 546 | 82.23% | 79.82% |
| | III.trna7 | iMet | Ile | AAT | 1 | 2 | 61.2 | 1 | 0.15% | 0.15% |
| *C. briggsae* | X.trna158 | Arg | His | ATG | 1 | 7 | 39.2 | 1 | 0.15% | 0.14% |
| | III.trna33 | Asn | Lys | TTT | 1 | 6 | 73.5 | 1 | 0.14% | 0.14% |
| | IV.trna54 | Asn, Trp, Tyr | Sup, Trp | TCA, CCA | 5 | 12 | 26.1 | 339 | 56.69% | 47.15% |
| | V.trna30 | Asn | His | GTG | 1 | 5 | 46.8 | 2 | 0.33% | 0.28% |
| | IV.trna1 | Asp | Gly | GCC | 1 | 14 | 46.8 | 1 | 0.17% | 0.14% |
| | X.trna47 | Gln | Sup | TTA | 1 | 5 | 45 | 2 | 0.28% | 0.28% |
| | II.trna110 | Glu | Arg, Undet | TCT, NNN | 1 | 8 | 50.55 | 18 | 2.80% | 2.50% |
| | II.trna38 | Glu | Sup | TCA | 1 | 6 | 64.6 | 1 | 0.14% | 0.14% |
| | I.trna134 | Gly | Arg | TCT | 1 | 4 | 71.7 | 1 | 0.14% | 0.14% |
| | I.trna49 | Gly | Glu | TTC | 1 | 9 | 67.2 | 1 | 0.14% | 0.14% |
| | II.trna81 | Gly | Ser, Undet | CGA, NNN | 1 | 4 | 47.55 | 10 | 1.41% | 1.39% |
| | V.trna134 | Gly | Ser | GCT | 1 | 10 | 77.2 | 1 | 0.14% | 0.14% |
| | X.trna117 | Gly | Arg | TCT | 1 | 7 | 45.9 | 1 | 0.15% | 0.14% |
| | X.trna133 | Ile | Val | TAC | 1 | 8 | 75.4 | 1 | 0.14% | 0.14% |

| | | | | | | | | | | |
|---|---|---|---|---|---|---|---|---|---|---|
| | II.trna90 | Lys | Sup | CTA | 1 | 5 | 86 | 21 | 2.95% | 2.92% |
| | V.trna15 | Pro | Asn | ATT | 1 | 2 | 56.3 | 1 | 0.15% | 0.14% |
| | I.trna101 | Thr | Ile | AAT | 1 | 4 | 84.4 | 1 | 0.14% | 0.14% |
| | I.trna129 | Thr | Ile | AAT | 1 | 4 | 83 | 2 | 0.28% | 0.28% |
| | I.trna47 | Thr | Ile | AAT | 1 | 4 | 85 | 2 | 0.28% | 0.28% |
| | I.trna65 | Thr | Lys | TTT | 2 | 7 | 72.4 | 4 | 0.57% | 0.56% |
| | II.trna44 | Thr | Ser | AGA | 1 | 3 | 85.3 | 2 | 0.28% | 0.28% |
| | II.trna67 | Thr | Ile | AAT | 1 | 6 | 84.6 | 12 | 1.74% | 1.67% |
| | II.trna86 | Thr | Ile | AAT | 1 | 2 | 83 | 1 | 0.14% | 0.14% |
| | II.trna89 | Thr | Ile | AAT | 2 | 6 | 86.75 | 5 | 0.70% | 0.70% |
| | III.trna113 | Thr | Ile | AAT | 1 | 4 | 79.1 | 5 | 0.72% | 0.70% |
| | III.trna12 | Thr | Ile | AAT | 1 | 4 | 79.1 | 1 | 0.14% | 0.14% |
| | III.trna9 | Thr | Ile | AAT | 1 | 3 | 84.4 | 1 | 0.14% | 0.14% |
| | X.trna105 | Thr | Ile | AAT | 1 | 4 | 79.6 | 1 | 0.14% | 0.14% |
| | X.trna123 | Thr | Tyr | GTA | 1 | 6 | 60.9 | 1 | 0.14% | 0.14% |
| | IV.trna88 | Trp | Sup | TCA | 1 | 4 | 40.3 | 4 | 0.63% | 0.56% |
| | IV.trna89 | Trp | Gly | TCC | 1 | 4 | 42.6 | 1 | 0.15% | 0.14% |
| | I.trna54 | iMet | Leu | CAA | 1 | 9 | 61.2 | 1 | 0.14% | 0.14% |
| | III.trna106 | iMet | Trp | CCA | 1 | 5 | 43 | 18 | 2.53% | 2.50% |
| *C. tropicalis* | II.trna100 | Gly | Arg | TCG | 1 | 2 | 71.7 | 2 | 0.33% | 0.32% |
| | II.trna111 | Gly | Ser | GCT | 1 | 6 | 77.2 | 1 | 0.16% | 0.16% |
| | I.trna54 | Lys | Sup | TTA | 1 | 3 | 72.5 | 3 | 0.48% | 0.48% |
| | V.trna80 | Met | Ile | TAT | 1 | 4 | 77 | 1 | 0.16% | 0.16% |
| | II.trna36 | Thr | Lys | TTT | 1 | 4 | 59.2 | 2 | 0.32% | 0.32% |
| | II.trna76 | Thr | Ile | AAT | 1 | 2 | 85 | 1 | 0.16% | 0.16% |
| | I.trna19 | Trp | Arg | ACG | 1 | 3 | 39.1 | 1 | 0.16% | 0.16% |
