## Supplemental Figures for "Allelic variation at tRNA genes in three nematode species indicates mutation load despite strong purifying selection"

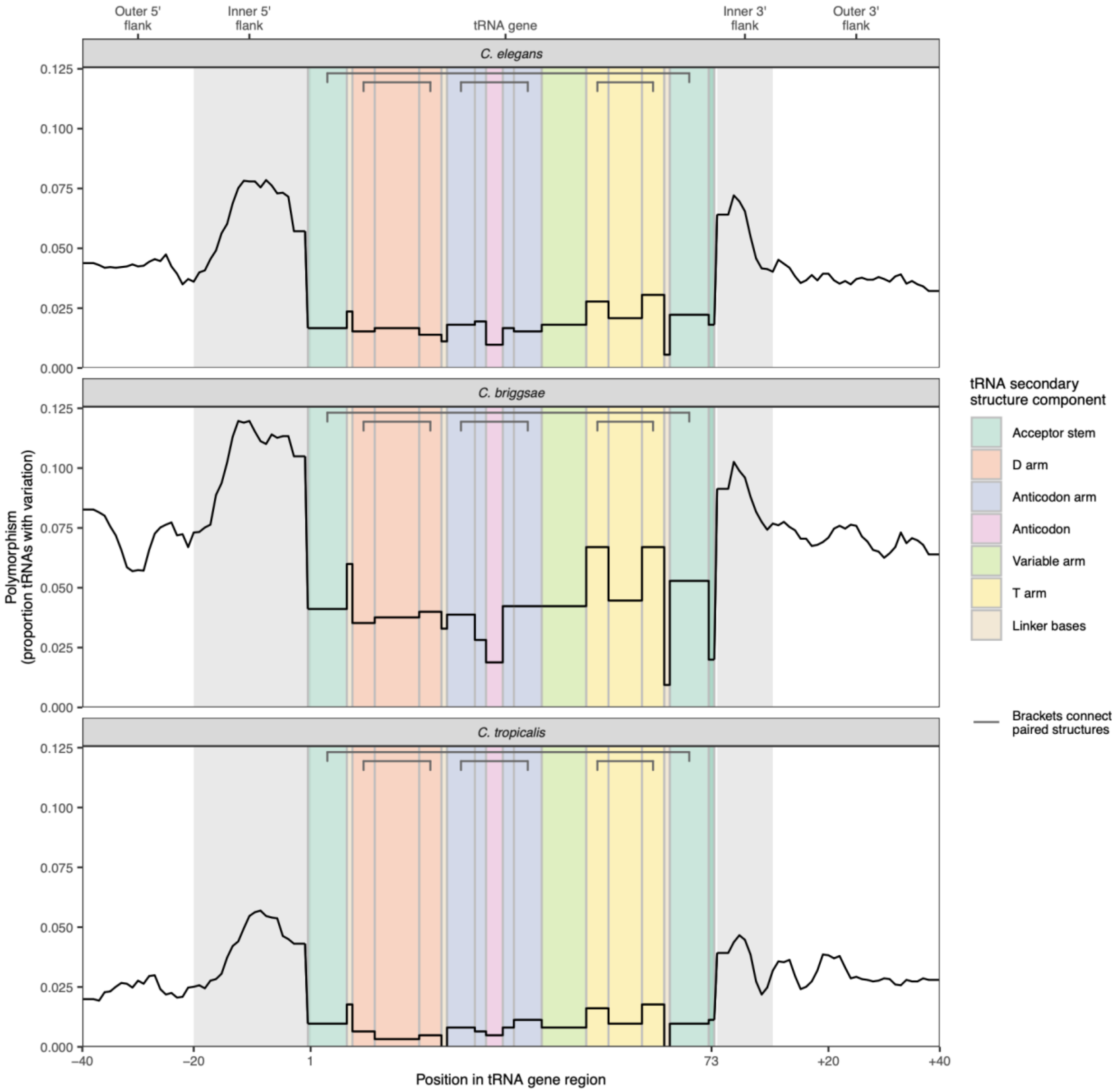


**Figure S1.** High rate of mutation in tRNA flanking regions in *Caenorhabditis* nematodes. As in **Figure 1C** but for all three species (sub-plot titles). Linker bases are included here (omitted from Figure 1); these do not occur in all tRNAs and therefore estimates for them are unreliable.

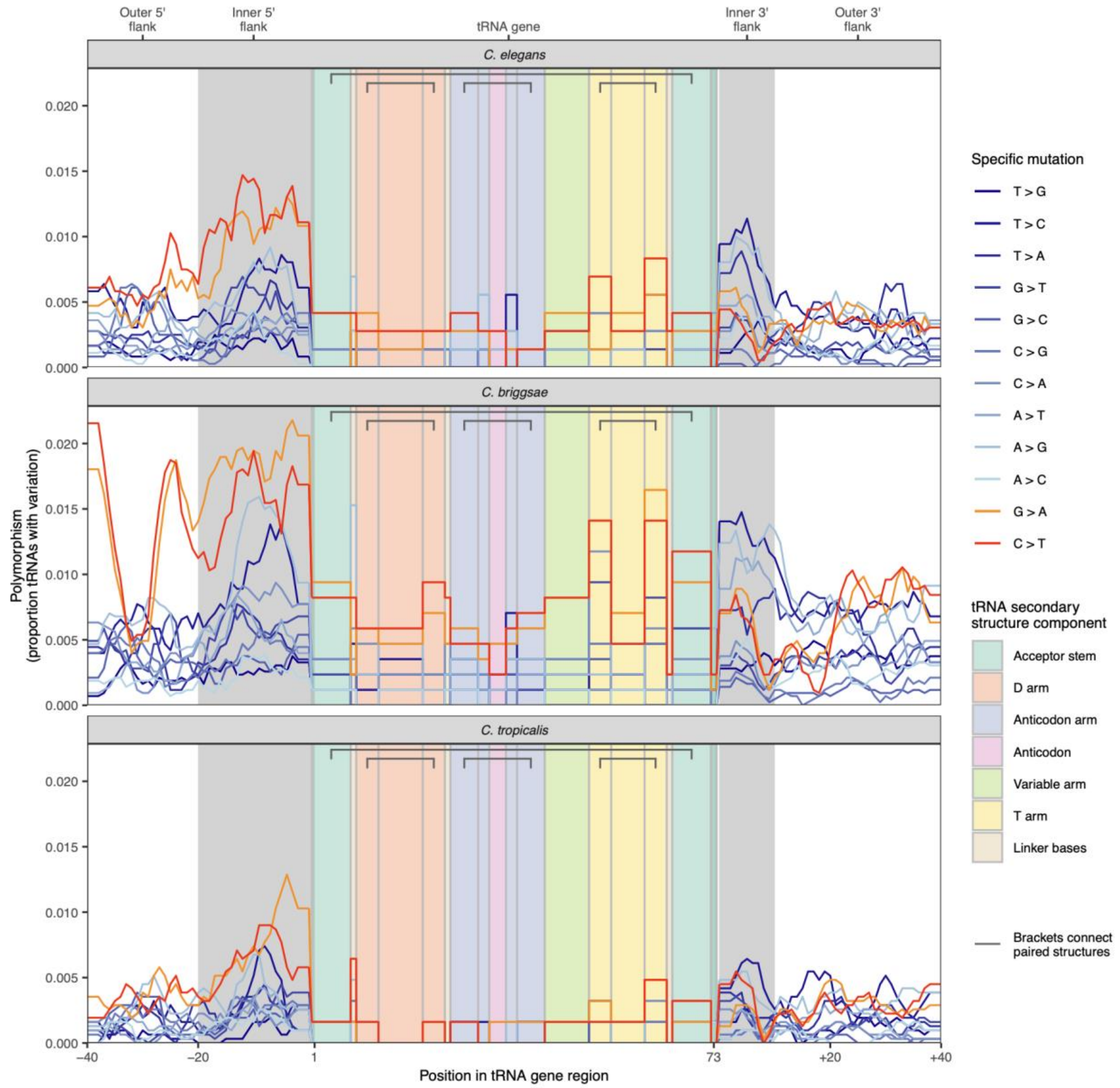


**Figure S2.** Signature of transcription-associated mutagenesis in *Caenorhabditis.* As in **Figure 1C,** but each class of SNV mutation is plotted separately in its own color, with TAM-characterizing mutations in orange and red. Linker bases are included here (omitted from Figure 1); these do not occur in all tRNAs and therefore estimates for them are unreliable.

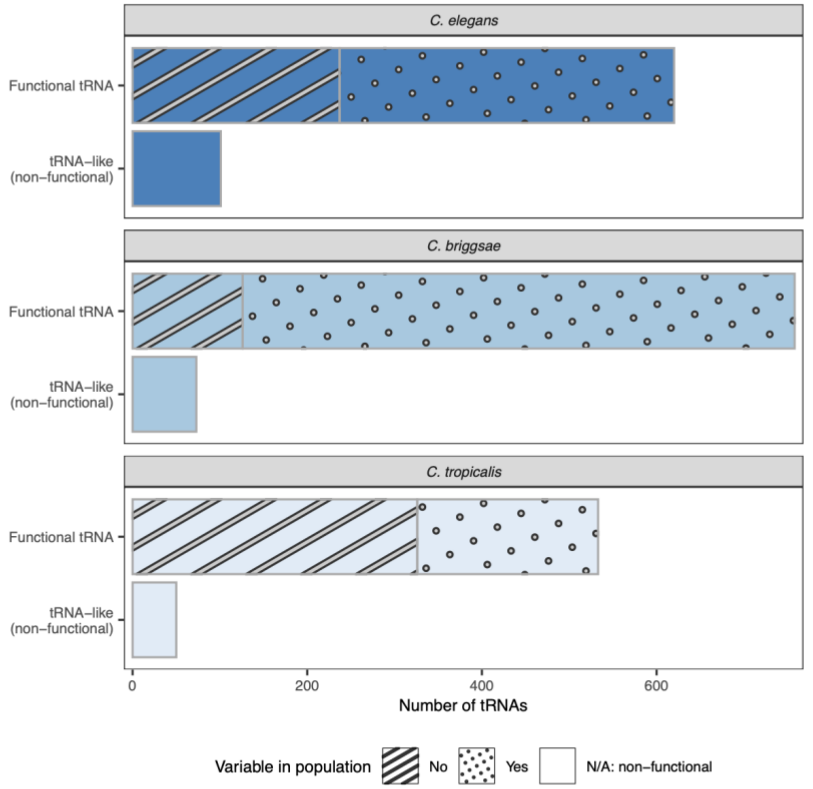


**Figure S3.** Total numbers of tRNA genes and tRNA-like sequences (pseudo genes) across *Caenorhabditis* species, subdivided by the number of genes with multiple alleles in the population (variable) vs fixed in the population (not variable).

(Related to **Figure 2**)

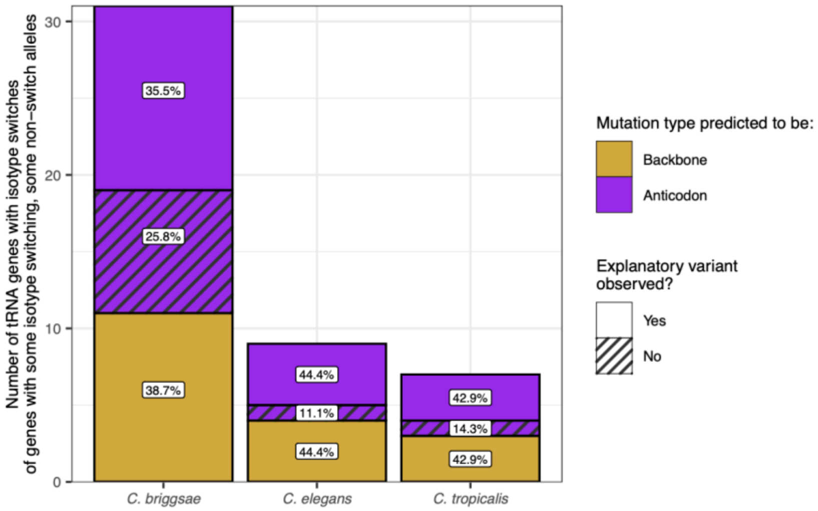


**Figure S4.** Characterization of variable isotype switch alleles by whether we can independently from tRNAscan-SE identify mutations likely underlying the isotype switch from the genetic variants. The proportion of each bar (per species) is shown on the plot, with the top and bottom bars summing to the total proportion of tRNAscan-SE called isotype switches for which we see independent mutational evidence. This is only for the 'some switches, some not' category (purple in **Figure 3A**) because if the isotype switch is fixed there is no variation in the population hence we cannot observe a variant causing the isotype switch. An explanatory variant not being observed could occur due to several factors including that the variant was an INDEL, which our methods cannot accurately map to structures (so they are included in allelic diversity but not per-variant analyses) ahd that the tRNA gene sequence was not appropriately broken down into secondary structure components and so we cannot analyze specific variants for that tRNA.

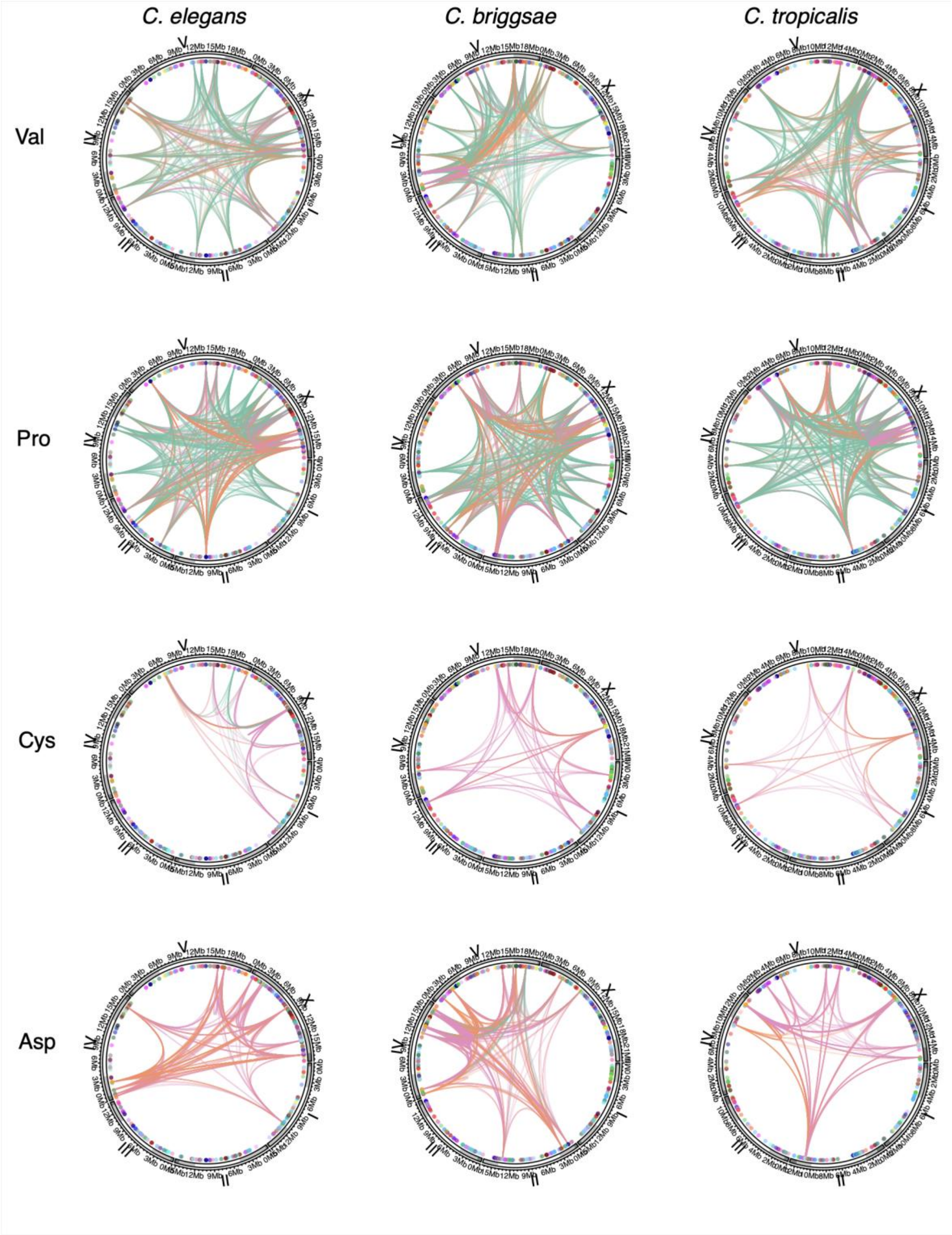


**Figure S5.** Distribution of tRNA genes in reference genomes and connections between tRNAs of specific isotypes. As in **Figure 4B** (including colors, see that key). All tRNAs are shown as dots in all plots in their genomic position (circular layout) per species (columns). The connections between all tRNAs with a given isotype (rows) are plotted.

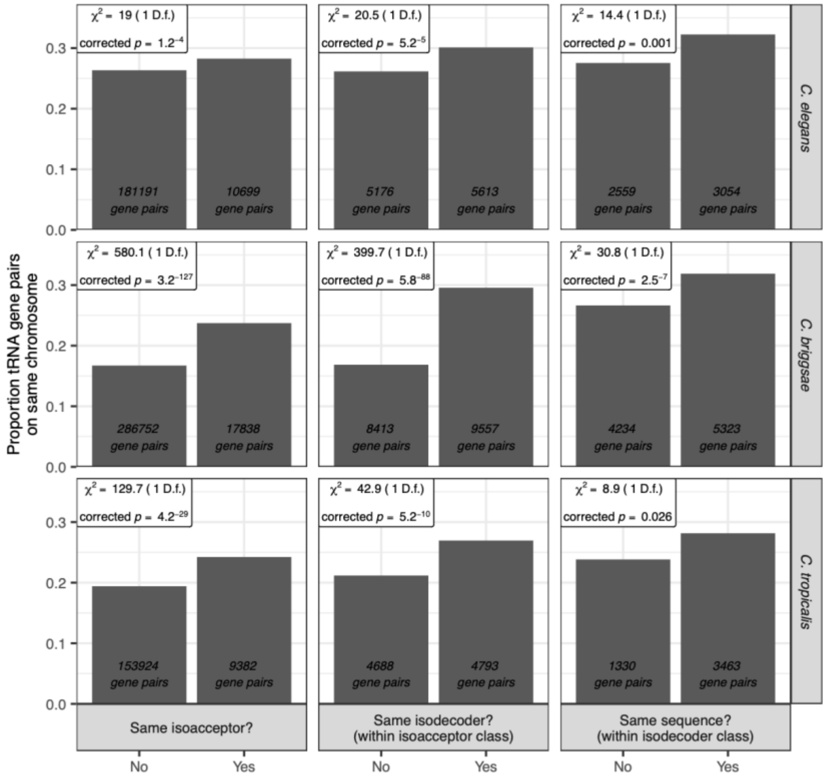


**Figure S6.** The proportion of tRNA gene pairs occurring on the same chromosome in the reference genome in three species (rows), subdivided by gene similarity or difference (columns). Statistical test results are shown for each comparison in each species; p-values shown are two-sided and Bonferroni corrected for multiple tests ($n$ = 9 tests; 3 species x 3 tests). All tRNA gene pairs between genes with at least one non-nonfunctional allele are included; $n$ pairs annotated on bars.

(Related to **Figure 4**)

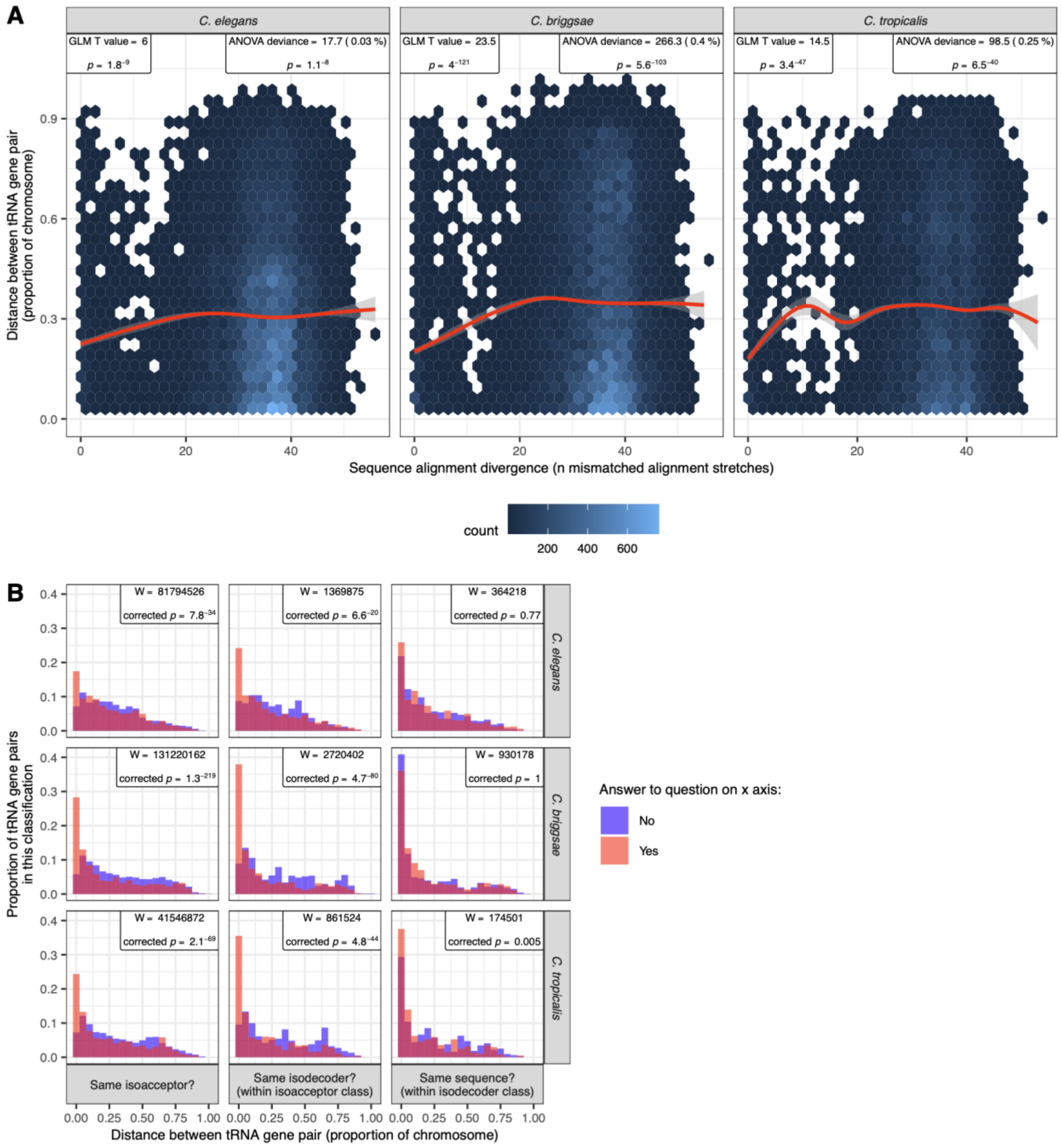


**Figure S7.** Relationships between tRNA gene sequence similarity and distance separating tRNA gene pairs in the genome. **A**, Quantitative relationship between sequence alignment divergence between tRNA pairs (x axis, see Methods) and distance between tRNA pairs (y axis), for all gene pairs occurring on the same chromosome. Statistical test results are shown from a generalized linear model (gamma log) modeling distance between genes as a function of chromosome, pseudogene status, and sequence divergence; ANOVA results show the deviance and percent deviance explained by sequence divergence from this GLM and GLM results are for the sequence divergence term. **B,** Distribution of distance between tRNA gene pairs when the pair does or does not share the provided characteristic (overlaid histograms; height of bar corresponds to proportion of the pairs in that category in that species). Statistical test results for Mann-Whitney tests are shown; p-values are two-sided and Bonferroni-corrected for multiple hypothesis testing (x 9: number of species x number of tests). Genes with all alleles classified as nonfunctional are excluded from this analysis (but included in A where pseudogene status can be included in statistical modeling).

(Related to **Figure 4**)
